# FBRSL1 couples ncPRC1.3 to CoREST to repress cell-cycle genes during human neuronal differentiation

**DOI:** 10.64898/2026.10.20.760506

**Authors:** Sanxiong Liu, Han-Wen Chang, Danny Reinberg

## Abstract

Most Polycomb complexes foster canonical gene repression, while noncanonical PRC1.3 (ncPRC1.3) unexpectedly activates transcription through its AUTS2 component. If and how ncPRC1.3 may also foster gene repression during development is unknown. Here, we show that in contrast to the impact of AUTS2 on ncPRC1.3, its paralog, FBRSL1, couples ncPRC1.3 to the CoREST/CTBP/LSD1 corepressor complex to mediate transcriptional repression. AUTS2 and FBRSL1 compete for binding to the RAWUL domain of the ncPRC1.3 subunit, PCGF3, and their isolated Atrophin-1 domains confer opposite transcriptional outputs. Corepressor interaction maps to a central Atrophin-1-containing region of FBRSL1, with CoREST and CTBP2 required for reporter repression. Patient-derived *FBRSL1* truncations abolish this repressive capacity; specifically, the p.Gln163* variant retains PCGF3 binding, but loses corepressor association. In human embryonic stem cells carrying a homozygous p.Gln163* knock-in, cortical differentiation displays persistent cell-cycle gene expression and deficient neuronal gene induction. At the progenitor stage, chromatin occupancy of CoREST, CTBP2, PCGF3, and RING1B is lost and accessibility increases at cell-cycle promoters that subsequently fail to be repressed. Thus, FBRSL1 redirects ncPRC1.3 function by equipping the complex with a corepressor-linked repressive arm that restrains proliferative programs at the progenitor-to-neuron transition.

**Highlights:**

- AUTS2 and FBRSL1 compete for PCGF3 leading to opposite ncPRC1.3 transcriptional outputs.
- FBRSL1 recruits CoREST/CTBP/LSD1 via its Atrophin-1 domain to repress transcription.
- Patient FBRSL1 truncations retain PCGF3 binding but lose corepressors and repression.
- BRSL1 truncation de-represses cell-cycle genes and derails neuronal differentiation.

**eTOC Blurb:** Liu et al. show that FBRSL1, a paralog of the Polycomb-associated activator AUTS2, links noncanonical PRC1.3 to the CoREST/CTBP/LSD1 corepressors. A patient-derived FBRSL1 truncation disrupts corepressor engagement while retaining ncPRC1.3 binding, causing persistent cell-cycle gene expression and impaired neuronal induction at the progenitor-to-neuron transition.

## Introduction

Most Polycomb group (PcG) complexes are canonical chromatin repressors that establish and maintain silent transcriptional states during development. In mammals, Polycomb repressive complex 1 (PRC1) comprises a diverse family of assemblies organized around a catalytic RING1A or RING1B core paired with one of six Polycomb group RING finger homologs (PCGF1–6, Gao et al. 2012). While canonical PRC1 complexes (incorporating PCGF2 or PCGF4) rely on chromobox (CBX) proteins to recognize the chromatin constituent, trimethylated lysine 27 of histone H3 (H3K27me3), noncanonical variant complexes (ncPRC1; containing PCGF1, PCGF3, PCGF5, or PCGF6) assemble with RYBP or YAF2 and independently catalyze histone H2A lysine 119 monoubiquitination (H2AK119ub1) (Gao et al. 2012; Blackledge et al. 2014; Fursova et al. 2019). The C-terminal RAWUL domain of PCGF proteins serves as a modular docking platform for lineage-specific accessory subunits, exemplified by BCOR recruitment to PCGF1 and PCGF3 (Junco et al. 2013). These accessory factors alter chromatin engagement and enzymatic output, allowing noncanonical assemblies to occupy transcriptionally active loci or function independently of RING1A/B catalytic activity (Scelfo et al. 2019). Although ncPRC1.3 and ncPRC1.5 complexes initiate Polycomb domain formation and gene silencing during X-chromosome inactivation(Almeida et al. 2017), how their regulatory activities are tailored by alternative accessory subunits across developmental lineages remains an open question.

The noncanonical assembly containing AUTS2 provides a striking precedent for how an accessory factor can reverse Polycomb function. When incorporated into ncPRC1.3 or ncPRC1.5, AUTS2 recruits casein kinase 2 (CK2), which phosphorylates RING1B to inhibit its H2A ubiquitin ligase activity, while simultaneously recruiting the transcription coactivator, histone acetyltransferase p300, via an alternating histidine-X (HX) repeat motif, converting ncPRC1 into a transcriptional activator (Gao et al. 2014; Liu et al. 2021). Nuclear respiratory factor 1 (NRF1) targets AUTS2-ncPRC1.3 to specific neuronal promoters to activate neurodevelopmental programs, and disrupting this pathway impairs motor neuron differentiation (Liu et al. 2021). AUTS2 can also cooperate with PRC2 to mediate gene repression in distinct progenitor contexts (Shimaoka et al. 2025). AUTS2 belongs to an ancient gene family that includes two vertebrate paralogs, FBRS and FBRSL1 (Sellers et al. 2020). FBRSL1 was originally identified alongside AUTS2 in PCGF3 and PCGF5 affinity purifications (Gao et al. 2012), and competitive association of AUTS2 and FBRSL1 for the PCGF3 scaffold has been proposed (Liu et al. 2021). However, whether FBRSL1 equips ncPRC1.3 with an activating output analogous to AUTS2, confers a distinct repressive activity, or acts neutrally has remained unknown.

Human genetics and functional modeling underscore the necessity of defining FBRSL1 function during development. *De novo* heterozygous truncating mutations in *FBRSL1* cause a severe neurodevelopmental syndrome characterized by microcephaly, intellectual disability, postnatal growth restriction, and craniofacial, skeletal, and cardiac malformations, displaying partial phenotypic overlap with AUTS2 syndrome (Ufartes et al. 2020; Pauli et al. 2021; Bukvic et al. 2024; Xu et al. 2026). *FBRSL1* is expressed in neural progenitor cells (Xu et al. 2026), and patient-derived mutant transcripts escape nonsense-mediated decay (Ufartes et al. 2020). FBRSL1 generates multiple alternative transcripts, and studies in *Xenopus* have linked short N-terminal isoforms to cranial neural crest development and alternative *mdm2* splicing via association with the splicing factor SF3B1 (Gerstner et al. 2025). Furthermore, FBRSL1 has been reported to associate with YY1 to support the expression of the histone acetyltransferases *BRPF1* and *KAT6A* (Kastens et al. 2025). Despite these phenotypic connections, the biochemical effector machinery engaged by full-length (FL) FBRSL1 within the noncanonical PRC1 scaffold, and how patient-derived truncations alter its chromatin-regulatory function during human neurogenesis, remain unresolved.

The neurodevelopmental manifestations of *FBRSL1* mutations highlight the fundamental process of cortical neurogenesis, wherein neural progenitors must exit the cell cycle to commit to neuronal lineages (Ohnuma and Harris 2003; Götz and Huttner 2005; Dehay and Kennedy 2007). This transition requires the coordinated silencing of proliferative gene programs. Classic transcriptional corepressor complexes containing CoREST (RCOR1), CTBP1/2, the histone demethylase LSD1 (KDM1A), and HDAC1/2 coordinate the demethylation of histone H3 lysine 4 with histone deacetylation (Y. Shi et al. 2003; Y.-J. Shi et al. 2005; Lee et al. 2005). Although CoREST complexes were originally characterized as REST-dependent corepressors that maintain neuronal gene silence in non-neuronal cells (Andrés et al. 1999; Ballas et al. 2005), their developmental output depends entirely on the recruiting partner: while LSD1 and RCOR2 can sustain neural progenitor self-renewal (Sun et al. 2010; Wang et al. 2016), CoREST complexes recruited by factors such as INSM1 enforce cell-cycle exit and promote cortical differentiation (Fuentes et al. 2012; Monaghan et al. 2017). Although PcG proteins are known to regulate neural stem cell fate (Corley and Kroll 2015), whether a noncanonical Polycomb accessory subunit can directly couple the PRC1 scaffold to corepressor machinery to extinguish proliferative programs during neurogenesis has not been established.

Here, we identify FBRSL1 as an adaptor that couples ncPRC1.3 to the CoREST/CTBP/LSD1 corepressor complex, establishing a repressive counterpart to AUTS2. We demonstrate that AUTS2 and FBRSL1 compete for the PCGF3 scaffold, with their homologous Atrophin-1 regions encoding opposite transcriptional outputs. This modular architecture is selectively disrupted by patient-derived truncating mutations, which preserve Polycomb docking but sever corepressor recruitment. In human cortical differentiation models, this uncoupling leads to persistent cell-cycle gene expression and impaired neuronal induction, uncovering a chromatin-based mechanism that restrains progenitor proliferation to enable human neurogenesis.

## Results

### FBRSL1 and AUTS2 incorporate individually into ncPRC1.3, compete for binding to the PCGF3 RAWUL domain, and recruit distinct effectors for opposite transcriptional regulation

To investigate how related accessory proteins contribute to the biochemical diversity and transcriptional output of noncanonical Polycomb repressive complex 1.3 (ncPRC1.3), we examined the AUTS2 family of proteins in the context of the PCGF3 scaffold. A re-analysis of our earlier proteomic survey of human PRC1 complexes (Gao et al. 2012) revealed that PCGF3, the defining component of ncPRC1.3, co-purifies from HEK293T-REx nuclear extracts with the core catalytic subunits RING1A and RING1B, the auxiliary factors RYBP and YAF2, casein kinase 2 (CK2), and all three members of the AUTS2 family: AUTS2, FBRS, and FBRSL1 (Figure 1A). AUTS2, FBRS, and FBRSL1 share a conserved domain architecture comprising an N-terminal PAT1-homology domain, an Atrophin-1-like region containing an alternating histidine-X (HX) repeat motif, and a central AUTS2 domain (Figure 1B, (Sellers et al. 2020)). Despite their shared ability to incorporate into ncPRC1.3, only AUTS2 has a defined function in this complex through its recruitment of the acetyltransferase p300 and resultant transcription activation (Gao et al. 2014; Liu et al. 2021). Whether FBRS and FBRSL1 confer similar transcriptional activation, mediate repression, or act neutrally was unknown.

**Figure 1.**
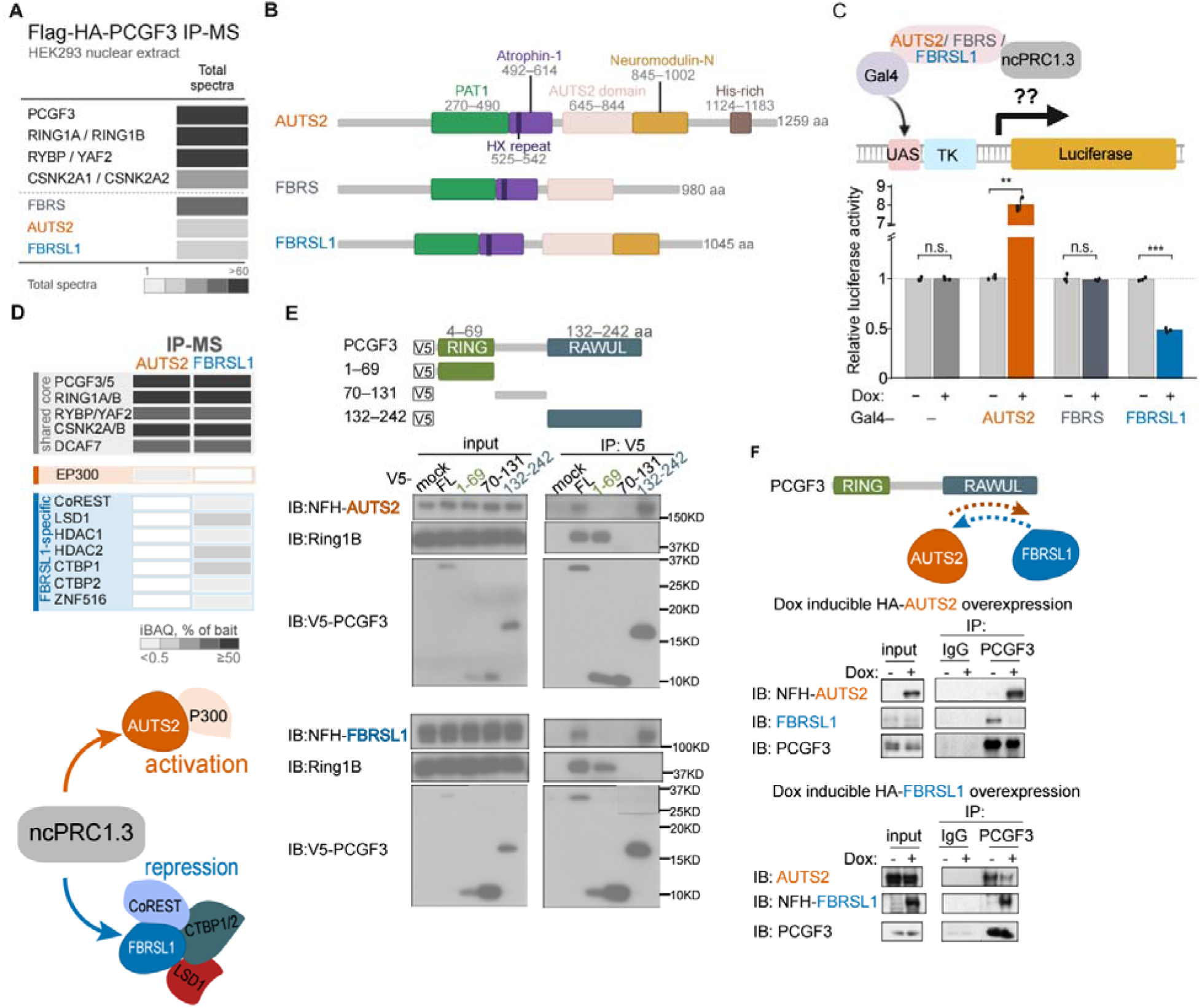
AUTS2 and FBRSL1 compete for PCGF3 binding and confer opposite transcriptional outputs on ncPRC1.3. **(A)** Selected proteins identified by immunoprecipitation–mass spectrometry (IP–MS) of Flag– HA-PCGF3 from HEK293T-REx nuclear extract, including noncanonical Polycomb repressive complex 1.3 (ncPRC1.3) components, casein kinase 2 subunits and AUTS2-family proteins (Gao et al., 2012). Shading indicates total spectral counts, in bins of 1–5, 6–15, 16–30, 31–60 and >60. **(B)** Domain and motif schematics of human AUTS2, FBRS and FBRSL1 protein. PAT1, Atrophin-1, HX repeat, AUTS2 domain, Neuromodulin-N, and His-rich annotations and amino-acid coordinates are indicated. **(C)** Top: schematic of the Gal4–UAS–TK–luciferase reporter assay. Doxycycline (Dox) induces the expression of the Gal4 DNA-binding domain (DBD), either alone or fused with AUTS2, FBRS or FBRSL1 in HEK293T-REx cells carrying a stably integrated 5×UAS–TK–luciferase reporter. Bottom: relative luciferase activity without (−) or with (+) Dox for 24 h. Luciferase activity was normalized to total protein. Bars show mean ± SD of n = 3 replicates, with dots for individual replicates. Statistics: Welch’s two-tailed t-test, +Dox versus −Dox for each construct, **p < 0.01; ***p < 0.001; n.s., not significant. **(D)** Top: comparative IP-MS of Flag-HA (NFH)-AUTS2 and NFH-FBRSL1 performed in HEK293T-REx nuclear extracts. Shading represents intensity-based absolute quantification (iBAQ), expressed as a percentage of the corresponding bait iBAQ, in bins of <0.5, 0.5–2, 2–10, 10–50 and ≥50%. White cells denote proteins not detected. Bottom: the accompanying schematic distinguishes the two ncPRC1.3 subcomplexes, with one harboring AUTS2 and p300, and the other FBRSL1 and CoREST/CTBP/LSD1 corepressors. **(E)** Top: V5-tagged FL PCGF3 (1–242 aa) and fragments thereof: 1–69, 70–131 and 132–242, encompassing the RING or RAWUL regions as indicated. Bottom: anti-V5 IP from NFH-AUTS2 cells (upper blots) or NFH-FBRSL1 cells (lower blots) transfected with the indicated V5-PCGF3 constructs or empty vector (mock). Blots were probed for the NFH-tagged paralog, RING1B and V5. **(F)** Top: schematic of AUTS2 and FBRSL1 competing for binding to the PCGF3 RAWUL domain. Bottom: IP of endogenous PCGF3, or IgG control, from cells with Dox-inducible NFH-AUTS2 (upper) or NFH-FBRSL1 (lower), with or without Dox.

To directly compare the transcriptional regulatory potential of these three paralogs, we tethered the FL version of each protein to an integrated 5xUAS-TK–luciferase reporter in HEK293T-REx cells via the Gal4 DNA-binding domain (DBD) upon doxycycline (Dox) induction (Figure 1C). Tethered Gal4-AUTS2 stimulated reporter activity approximately eightfold, reproducing its established activating output (Gao et al. 2014; Liu et al. 2021). In sharp contrast, tethered Gal4-FBRSL1 repressed reporter activity by approximately 50%, whereas Gal4-FBRS and the Gal4 DBD alone elicited no significant transcriptional change (Figure 1C). Thus, AUTS2 and FBRSL1 execute opposite transcriptional activities when recruited to the same promoter context.

Because AUTS2-mediated activation is driven by p300 recruitment, we hypothesized that FBRSL1-mediated repression is executed through distinct, corepressor effector machinery. To test this possibility, we purified Flag-HA-tagged FBRSL1 from nuclear extracts using sequential phosphocellulose (P11) chromatography and Flag affinity purification (Figures S1A and S1B) and compared its associated proteins with those of Flag-HA-tagged AUTS2 (Gao et al. 2014) by mass spectrometry (IP–MS) (Figure 1D). Both baits recovered the common ncPRC1.3 core complex, including PCGF3, PCGF5, RING1A, RING1B, RYBP, YAF2, CK2 subunits, and DCAF7. However, their effector associations were mutually exclusive: p300 was recovered exclusively with AUTS2, whereas FBRSL1 specifically enriched classic corepressor components, including CoREST (RCOR1), the histone demethylase LSD1 (KDM1A), HDAC1, HDAC2, CTBP1, CTBP2, and the zinc-finger protein ZNF516 (Figure 1D).

We validated these MS identifications using independent biochemical approaches. Anti-Flag IP from cells expressing Dox-inducible Flag-HA-FBRSL1 confirmed robust co-precipitation of ncPRC1.3 core components (PCGF3, RING1B, RYBP) along with CoREST, LSD1, and CTBP1 exclusively upon Dox induction (Figure S1C). Reciprocal IPs of endogenous CTBP2, CoREST, RING1B, and PCGF3 from native nuclear extracts each successfully recovered endogenous FBRSL1 (Figure S1D). Furthermore, anti-Flag IP of tagged AUTS2 recovered PCGF3 but failed to detect CTBP2, despite abundant CTBP2 in the input extract (Figure S1E), demonstrating that corepressor interaction is a unique property of FBRSL1 and not a feature shared by other AUTS2-family members.

Because AUTS2 and FBRSL1 engage distinct effectors within a shared complex, we examined how both paralogs incorporate into ncPRC1.3 through PCGF3. We expressed V5-tagged FL PCGF3 or truncation fragments encompassing the N-terminal RING domain (residues 1–69), the central linker (residues 70–131), or the C-terminal RAWUL domain (residues 132–242) in cells expressing Flag-HA-tagged AUTS2 or FBRSL1. Anti-V5 IP revealed that while RING1B associated with FL PCGF3 and its RING fragment, both AUTS2 and FBRSL1 bound FL PCGF3 and the RAWUL domain, with neither paralog binding the RING or linker segments (Figure 1E).

Because AUTS2 and FBRSL1 bind the same PCGF3 RAWUL region, we tested whether they compete for incorporation into endogenous ncPRC1.3. We IP’ed endogenous PCGF3 from cells following Dox-induced overexpression of Flag-HA-AUTS2 or Flag-HA-FBRSL1 (Figure 1F). Overexpression of Flag-HA-AUTS2 displaced endogenous FBRSL1 from PCGF3, and conversely, overexpression of Flag-HA-FBRSL1 displaced endogenous AUTS2 from PCGF3, without altering the total recovery of PCGF3 (Figure 1F). These reciprocal displacement assays demonstrate that AUTS2 and FBRSL1 compete for the PCGF3 RAWUL domain to form two mutually exclusive ncPRC1.3 subcomplexes of opposite transcriptional regulation.

### FBRSL1 co-assembles with ncPRC1.3 and corepressors into a holocomplex required for transcriptional repression

Although FBRSL1 co-precipitates with both ncPRC1.3 core components and corepressors, these pairwise interactions could conceivably reflect two independent cellular pools of FBRSL1: one engaged with ncPRC1.3 and another with CoREST/CTBP/LSD1. To determine whether these factors assemble into a single, unified holocomplex, we subjected Flag-affinity-purified FBRSL1 complexes to Superose 6 size-exclusion chromatography (Figure 2A). Immunoblotting across the fractionation profile revealed that FBRSL1, CoREST, and CTBP1 co-eluted with the ncPRC1.3 subunits PCGF3, RING1B, and RYBP in the same high-molecular-weight fractions (fractions B4–B6) (Figure 2A). FBRSL1 and ncPRC1.3 core subunits were also detected in fractions flanking this peak that lacked detectable CoREST and CTBP1, indicating that FBRSL1-ncPRC1.3 can also exist independently of these corepressors—either as core complexes alone or in association with alternative partners. Importantly, the co-fractionation in fractions B4–B6 demonstrates that FBRSL1, ncPRC1.3, and corepressors co-assemble into a stable, high-molecular-weight macromolecular holocomplex.

**Figure 2.**
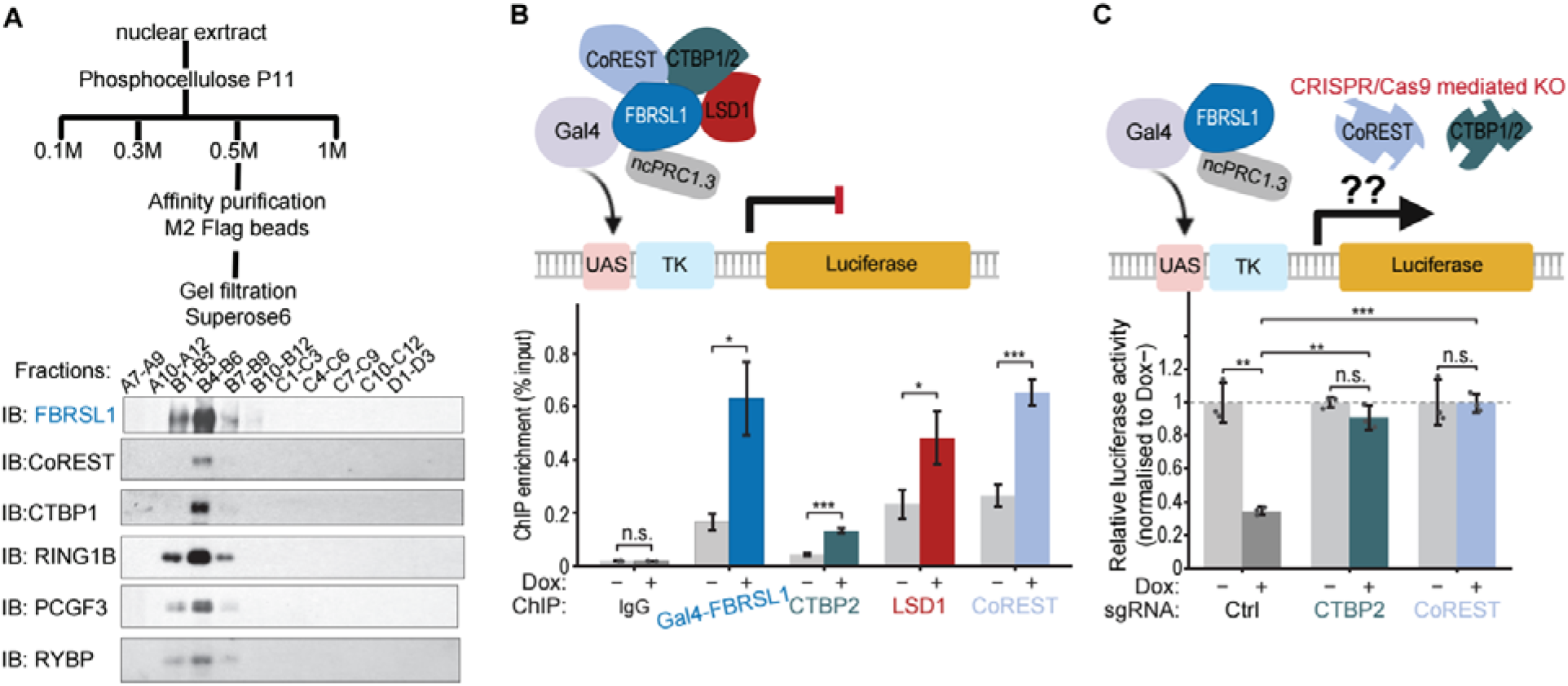
CoREST/CTBP/LSD1 corepressors assemble with FBRSL1–ncPRC1.3 and are required for transcriptional repression. **(A)** Top: biochemical purification scheme. Nuclear extract from HEK293T-REx NFH-FBRSL1 cells was fractionated on phosphocellulose P11 by step elution at 0.1, 0.3, 0.5 and 1.0 M KCl. The 0.5 M fraction was affinity-purified on anti-Flag M2 beads, eluted with Flag peptide and resolved by Superose 6 gel filtration. Bottom: immunoblots of the indicated fraction pools for FBRSL1, CoREST, CTBP1, RING1B, PCGF3, and RYBP. **(B)** Schematic of GAL4-FBRSL1-mediated corepressor recruitment to the UAS–TK–luciferase reporter (top) and chromatin immunoprecipitation followed by quantitative PCR (ChIP–qPCR) at the reporter regulatory region (bottom). Enrichment of GAL4-FBRSL1, CTBP2, LSD1 and CoREST was measured without (−) or with (+) doxycycline (Dox); IgG is the negative control. Bars show mean ± SD of n = 3 replicates. Statistics: two-tailed unpaired t-test, +Dox versus −Dox. *p < 0.05; ***p < 0.001; n.s., not significant. **(C)** Experimental schematic (top) and luciferase activity (bottom) in GAL4-FBRSL1 reporter cells carrying control sgRNA (Ctrl) or sgRNAs targeting CTBP2 or CoREST. Values are normalized to the −Dox mean of each line. Bars show mean ± SD of n = 3 replicates. Statistics: two-tailed unpaired t-test. **p < 0.01; ***p < 0.001; n.s., not significant. Knockout efficiency and Gal4-FBRSL1 induction are shown in Figure S2A.

We next tested whether tethered FBRSL1 physically recruits these corepressors to a chromatin template. Chromatin immunoprecipitation followed by quantitative PCR (ChIP–qPCR) at the integrated UAS–TK promoter showed that Dox-induced Gal4-FBRSL1 binding led to a marked, simultaneous increase in local chromatin occupancy of CTBP2, LSD1, and CoREST, whereas the control IgG signal remained background-level and unchanged (Figure 2B). Thus, FBRSL1 directly recruits the assembly of corepressor machinery at target chromatin loci.

To establish whether the recruited corepressors are functionally required for FBRSL1-mediated repression, we disrupted *RCOR1* (CoREST) or *CTBP2* in the Gal4-FBRSL1 reporter line using CRISPR–Cas9. Immunoblotting confirmed efficient ablation of CoREST and CTBP2 while preserving Dox-inducible Gal4-FBRSL1 expression (Figure S2A). In control cells, Gal4-FBRSL1 induction reduced luciferase activity as expected. In contrast, depletion of either CoREST or CTBP2 completely abrogated transcriptional repression (Figure 2C). Therefore, CoREST and CTBP2 are obligatory functional effectors required for FBRSL1-mediated gene silencing.

### An Atrophin-1-containing region of FBRSL1 is modular, separable from PCGF3 binding, and sufficient for repression

To elucidate the molecular architecture governing FBRSL1 function, we performed domain-mapping experiments to identify the regions that interact with PCGF3 and the corepressor complex. We expressed HA-tagged FBRSL1 or three non-overlapping fragments spanning the N-terminal region (residues 1–163), the central region (residues 164–778), and the C-terminal region (residues 779–1045) in HEK293T cells (Figure 3A). IP with anti-HA antibodies revealed that the N-terminal 1–163 fragment bound PCGF3 but failed to co-precipitate LSD1, CoREST, or CTBP2. Conversely, the central 164–778 fragment robustly recovered LSD1, CoREST, and CTBP2, but completely lacked PCGF3 binding. The C-terminal fragment failed to bind any of these components (Figure 3A). Thus, FBRSL1 possesses a modular adaptor architecture in which N-terminal PCGF3 docking is physically decoupled from central corepressor recruitment.

**Figure 3.**
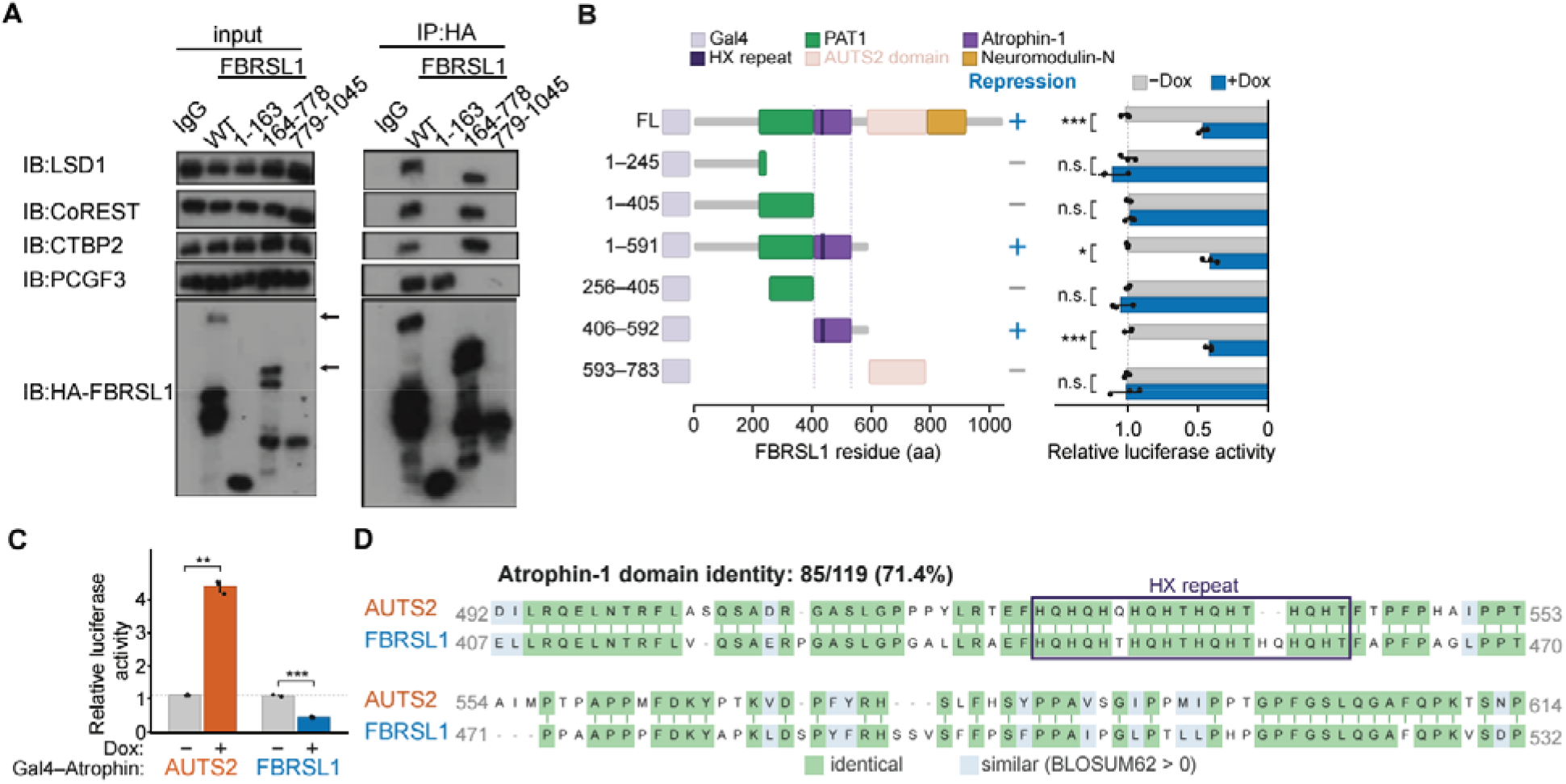
The Atrophin-1 domain of FBRSL1 engages the CoREST/CTBP/LSD1 complex to drive transcriptional repression. **(A)** Anti-HA IP of HA-tagged FL FBRSL1 (WT) or FBRSL1 fragments 1–163, 164–778, and 779– 1045, with IgG as the control, from HEK293T-REx cells. Blots were probed for LSD1, CoREST, CTBP2, PCGF3, and HA-FBRSL1. Arrows mark protein migrating at the predicted molecular weight. **(B)** GAL4-FBRSL1 construct schematics and corresponding reporter activities for FL FBRSL1 and fragments 1–245, 1–405, 1–591, 256–405, 406–592, and 593–783, without (−) or with (+) doxycycline (Dox). Dashed guides mark the Atrophin-1 core. The repression column summarizes significant repression (+) or no significant repression (−) in this assay. Bars show mean ± SD of n = 3 replicates, with dots for individual replicates. Statistics: Welch’s two-tailed t-test, +Dox versus −Dox. *p < 0.05; ***p < 0.001; n.s., not significant. **(C)** Relative luciferase activity of Gal4 fused to the Atrophin-1 domain of AUTS2 or FBRSL1, with or without Dox. Bars show mean ± SD of n = 3 replicates. Statistics: Welch’s two-tailed t-test. **p < 0.01; ***p < 0.001. **(D)** Pairwise alignment of the core Atrophin-1 domains of human AUTS2 (aa 492–614) and FBRSL1 (aa 407–532).

We next asked which region of FBRSL1 is sufficient to mediate transcriptional repression when tethered to DNA. We generated a series of Gal4 DBD fusions spanning truncated segments of FBRSL1 and tested them in the luciferase reporter assay (Figure 3B). While the N-terminal fragments (residues 1–245, 1–405) and an internal segment (residues 256–405) failed to repress luciferase expression, the 1–591 fragment and a central 406–592 fragment each conferred strong transcriptional repression comparable to FL FBRSL1 (Figure 3B). An adjacent downstream fragment (residues 593–783) exhibited no repressive activity. These functional mapping data define residues 406–592, which encompass the conserved Atrophin-1 homology domain, as a minimal segment sufficient for transcriptional repression.

Because the homologous region in AUTS2 drives transcriptional activation (Gao et al. 2014), we directly compared the transcriptional activity of the Atrophin-1-containing regions of both paralogs. We tested Gal4 fusions of the AUTS2 Atrophin-1 region (residues 405–717; (Gao et al. 2014)) alongside the FBRSL1 Atrophin-1 region (residues 406–592; Figure 3B) in the luciferase reporter assay. Remarkably, the two homologous domains produced opposite outputs: the Atrophin-1 domain of AUTS2 activated transcription approximately fourfold, whereas that of FBRSL1 repressed transcription by approximately 55% (Figure 3C). Notably, this functional divergence occurs despite profound sequence conservation: pairwise alignment of the core Atrophin-1 domains of AUTS2 (residues 492–614) and FBRSL1 (residues 407–532) revealed 71.4% sequence identity (85/119 identical residues), with near-perfect preservation of the HX repeat (Figure 3D). Therefore, despite high primary sequence similarity, the Atrophin-1-containing regions of AUTS2 and FBRSL1 have evolved opposite transcriptional activities, reflecting recruitment of p300 and of corepressor complexes, respectively.

### Patient-derived FBRSL1 truncations uncouple PCGF3 binding from corepressor recruitment and abolish repression

*De novo* heterozygous truncating mutations in *FBRSL1* cause a severe human syndromic disorder featuring microcephaly, intellectual disability, developmental delay, and craniofacial anomalies (Ufartes et al. 2020; Pauli et al. 2021; Xu et al. 2026; Bukvic et al. 2024). Mapping published pathogenic mutations onto the *FBRSL1* gene structure revealed that all five reported truncating alleles—including nonsense variants (c.332G>A, p.Trp111*; c.487C>T, p.Gln163*) and frameshift variants (c.371dupC, p.Cys125Leufs*7; c.380dup, p.Ala128Cysfs*5; c.581_603del, p.Gln196Profs*161)—introduce premature termination codons upstream of the sequence encoding the central Atrophin-1 domain (Figure 4A). Our domain-mapping data (Figure 3) predicted that these early truncations would yield truncated proteins that preserve N-terminal PCGF3 binding, but be devoid of corepressor-binding and the repressive Atrophin-1 domain.

**Figure 4.**
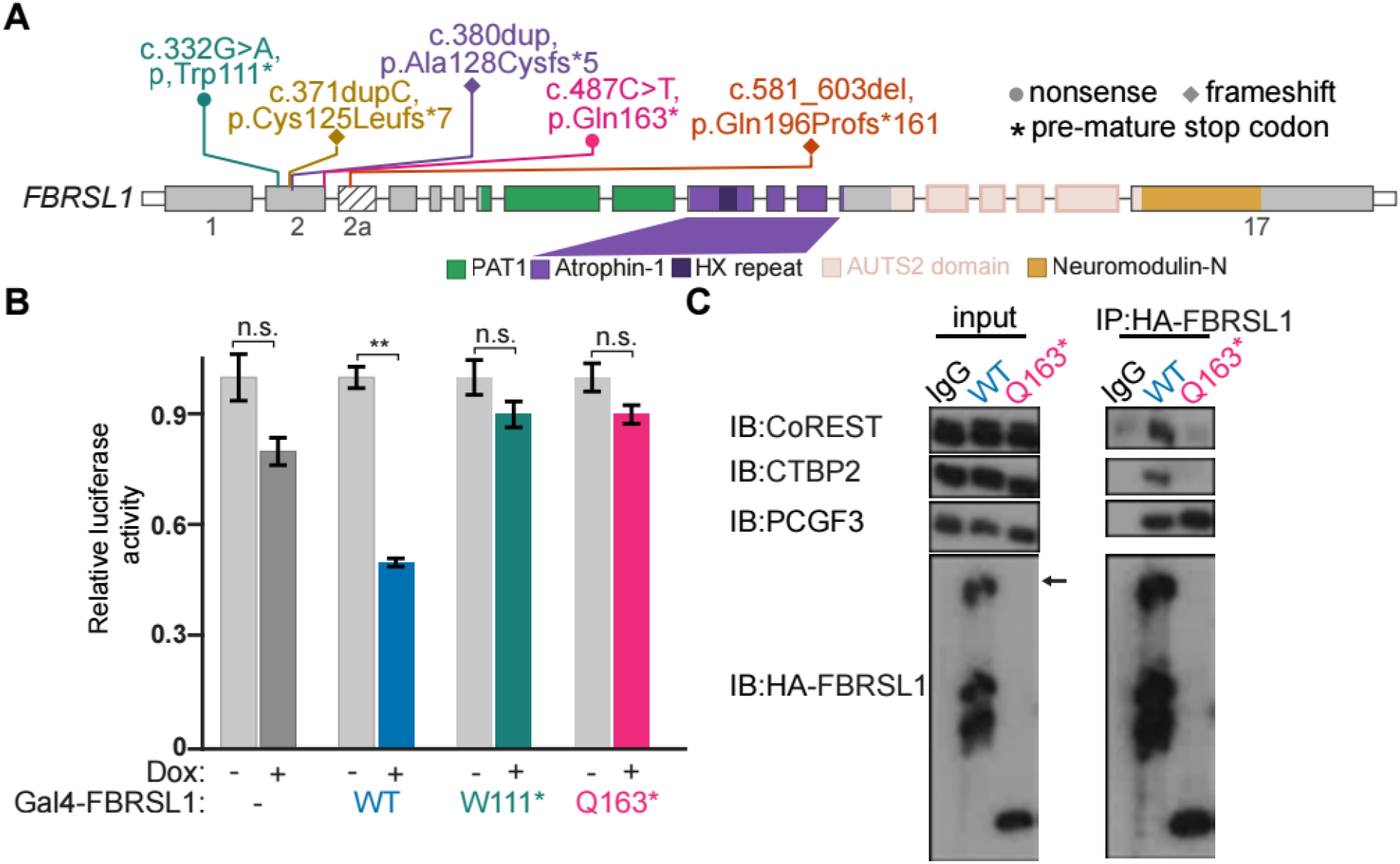
Patient-derived truncating mutations in FBRSL1 abolish its binding to the CoREST/CTBP complex and its repressive activity. **(A)** Schematic of the human *FBRSL1* gene showing five reported truncating variants with coding-DNA changes, predicted protein consequences, and domain positions. Circles indicate nonsense variants; diamonds indicate frameshift variants; asterisks indicate premature termination codons. **(B)** Gal4–UAS–TK–luciferase assay performed with the Gal4 DBD alone (−), or with GAL4 fusions with either WT FBRSL1, or the patient variants p.Trp111* and p.Gln163*, with or without Dox. Values are normalized to the −Dox mean of each construct. Bars show mean ± SD of n = 3 replicates. Statistics: Welch’s two-tailed t-test, +Dox versus −Dox. **p < 0.01; n.s., not significant. **(C)** Anti-HA IP of HA-tagged FBRSL1 WT or p.Gln163*, with IgG as the control, from HEK293T-REx cells. Blots were probed for CoREST, CTBP2, PCGF3, and HA-FBRSL1. The arrow marks FL FBRSL1.

We tested this prediction functionally using the Gal4 reporter assay. As expected, wild-type (WT) Gal4-FBRSL1 repressed luciferase activity by approximately 50%, however neither the patient-derived p.Trp111* nor p.Gln163* truncation mutant produced significant repression, exhibiting instead luciferase levels indistinguishable from those of the inactive Gal4 DBD control (Figure 4B).

To test whether this loss of repression stems from selective uncoupling of corepressors, we performed anti-HA co-immunoprecipitation comparing WT FBRSL1 and the patient-derived p.Gln163* variant. Whereas WT FBRSL1 co-precipitated PCGF3, CoREST, and CTBP2, the p.Gln163* truncated protein lost association with CoREST and CTBP2, while retaining robust binding to PCGF3 (Figure 4C). This biochemical behavior precisely matches that of the engineered 1–163 fragment (Figure 3A). Thus, recurrent patient-derived mutations represent molecular separation-of-function alleles: they produce truncated FBRSL1 proteins that preserve ncPRC1.3 scaffold docking but are physically uncoupled from corepressor machinery, thereby abolishing transcriptional repression.

### FBRSL1 p.Gln163* hESCs sustain cell-cycle gene expression and fail to induce neuronal differentiation

Because individuals harboring *FBRSL1* truncations suffer from profound neurodevelopmental defects, we asked how the p.Gln163* separation-of-function allele affects human neural differentiation. We knocked-in the patient-derived homozygous c.487C>T (p.Gln163*) mutation into the endogenous *FBRSL1* locus of H9 human embryonic stem cells (hESCs). WT and isogenic p.Gln163* hESCs were subjected to directed cortical neuronal differentiation using dual SMAD inhibition (Chambers et al. 2009; Edri et al. 2015), and transcriptomes were profiled by RNA-seq across five developmental stages: day 0 (D0, hESC), D4 (neuroectoderm), D9 (neural stem cell, NSC), D13 (neural progenitor cell, NPC), and D17 (cortical neurons) (Figure 5A). Consistent with reports that patient-derived FBRSL1 truncations escape nonsense-mediated decay (Ufartes et al. 2020), *FBRSL1* mRNA levels in mutant cells were comparable to those of WT throughout the time course of differentiation (Figure S3A).

**Figure 5.**
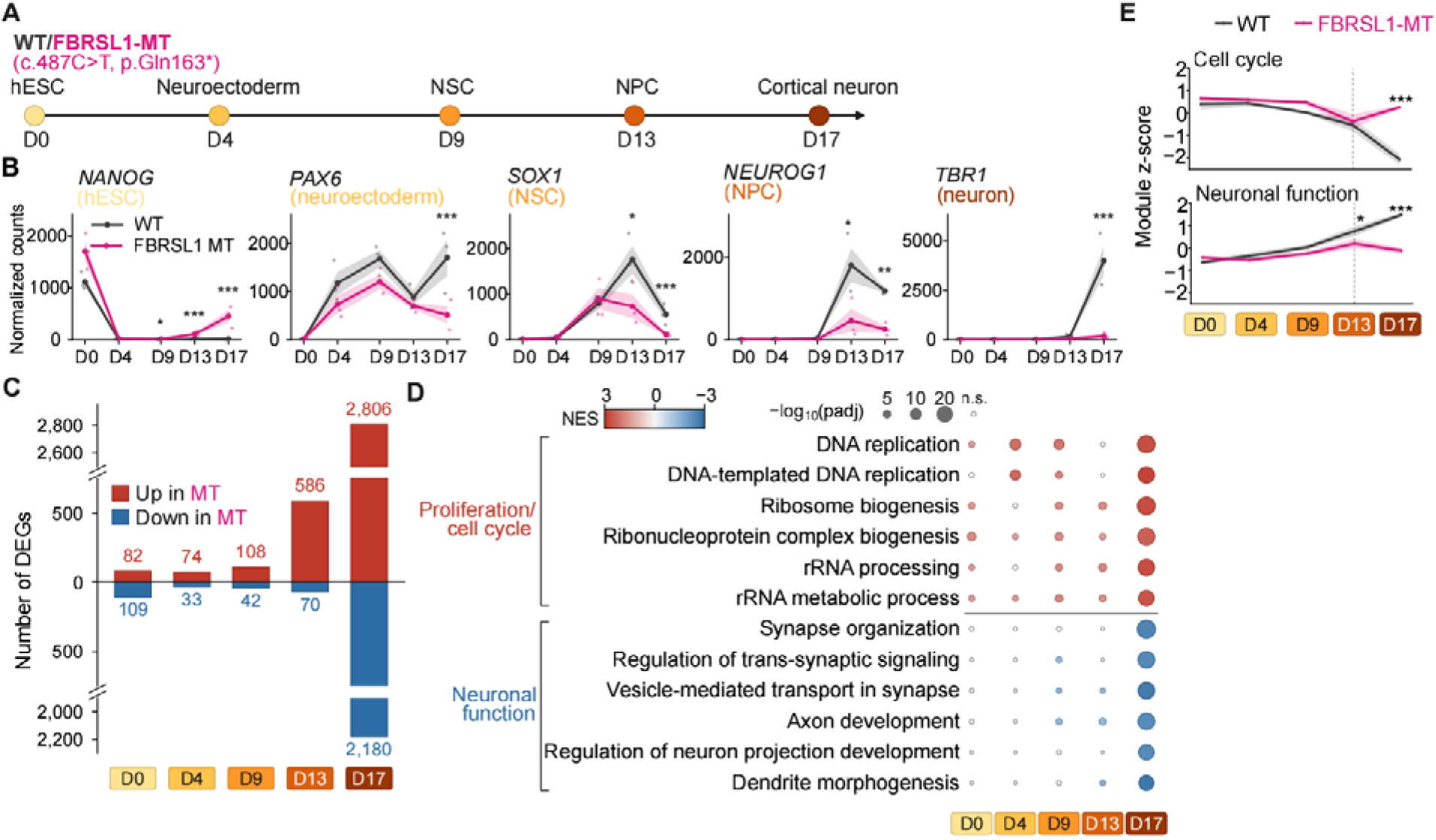
The FBRSL1 p.Gln163* mutation disrupts cell-cycle and neuronal gene expression programs during cortical differentiation of human ESCs. **(A)** Schematic of cortical neuron differentiation from WT H9 hESCs and from an isogenic hESC line, FBRSL1 mutant (FBRSL1-MT), carrying the homozygous patient-derived knock-in *FBRSL1* c.487C>T (p.Gln163*) at the endogenous locus. **(B)** Expression trajectories of stage-specific markers in WT and FBRSL1-MT cells across differentiation (D0–D17): NANOG (hESC), PAX6 (neuroectoderm), SOX1 (NSC), NEUROG1 (NPC) and TBR1 (cortical neuron). Values are DESeq2-normalized counts. Lines show the mean, shading shows ± SEM, and dots show individual replicates (n = 3). **(C)** Numbers of differentially expressed genes (DEGs) between FBRSL1-MT and WT at each time point (DESeq2; adjusted p < 0.05 and |log FC| ≥ 1). **(D)** Gene set enrichment analysis (GSEA) of GO Biological Process terms comparing FBRSL1-MT with WT at each time point. Genes were ranked by the DESeq2 Wald statistic and analyzed with fgsea. The six positively and six negatively enriched terms with the lowest adjusted p value across days are shown. **(E)** Expression trajectories of a cell-cycle module and a neuronal-function module containing genes that are significantly dysregulated in the FBRSL1 MT. Each gene was z-scored across all the libraries, and the module score is the mean z-score of all the genes in the module. Lines show the mean and shading shows ± SEM (n = 3). Statistics: two-way ANOVA (genotype × day). *p < 0.05; ***p < 0.001. The dotted line marks D13.

We first assessed staged lineage progression by tracking canonical marker genes (Figure 5B). Pluripotency-associated *NANOG* was properly downregulated by D4, and early neural stem-cell markers *PAX6* and *SOX1* were induced by D9 in both WT and mutant cells, indicating that initial neural induction proceeds without major impairment in the mutant (Figure 5B). However, severe divergence occurred at subsequent stages: between D13 and D17, the intermediate progenitor marker *NEUROG1* and post-mitotic cortical marker *TBR1* were robustly induced in WT cells, but their induction was severely attenuated in p.Gln163* cells, with *TBR1* expression being virtually undetectable at D17 (Figure 5B). Concurrently, *SOX1* and *PAX6* induction was not sustained at normal levels in the mutant at D17.

Principal component analysis (PCA) of the 500 most variable genes mirrored this developmental trajectory: WT and mutant transcriptomes clustered closely at D0, D4, and D9, began to separate at D13, and diverged dramatically at D17 (Figure S3B). Quantification of differentially expressed genes (DEGs; DESeq2, adjusted p < 0.05, |log_2_FC| ≥ 1) demonstrated a marked, stage-specific escalation in transcriptomic disruption: differences were modest at D0 (191 DEGs), D4 (107 DEGs), and D9 (150 DEGs), expanded to 656 DEGs at D13 (586 upregulated, 70 downregulated), and culminated in 4,986 DEGs at D17 (2,806 upregulated, 2,180 downregulated) (Figure 5C).

Gene set enrichment analysis (GSEA) of the dysregulated programs revealed two distinct, opposing transcriptional signatures (Figure 5D). Genes upregulated in mutant cells were heavily enriched for terms associated with cell proliferation and DNA synthesis, including DNA replication, cell cycle, ribosome biogenesis, and rRNA metabolic processing. Conversely, genes downregulated in the mutant were strongly enriched for mature neuronal functional terms, including synapse organization, trans-synaptic signaling, vesicle transport, axon guidance, and dendrite morphogenesis (Figure 5D).

Because the transition from neural progenitors (D13) to cortical neurons (D17) obligatorily requires cells to exit the cell cycle and activate neurogenic programs, we analyzed the expression trajectories of cell-cycle and neuronal-function gene modules across differentiation (Figure 5E). Between D13 and D17, WT cells successfully silenced the cell-cycle module while robustly activating the neuronal-function module. In contrast, p.Gln163* cells failed to execute both transitions: the cell-cycle module remained abnormally elevated at D17, whereas the neuronal module failed to turn on (Figure 5E). Importantly, the elevated expression of cell-cycle genes in mutant cells compared to WT at D17 does not reflect *de novo* transcriptional hyper-activation, but rather a failure of normal gene repression—resulting in the de-repression of proliferative programs. Concurrently, the reduced expression of neuronal genes represents an impaired induction of differentiation programs rather than active repression. Thus, mutant cells exhibit an uncoordinated transcriptional state characterized by persistent cell-cycle gene expression and failure to mount neuronal differentiation programs.

An analysis of epigenetic regulator dynamics in WT cells provided a temporal and mechanistic rationale as to why the phenotype manifests precisely at the D13 progenitor stage: *FBRSL1* and *PCGF3* mRNA levels rose sharply from D9 to D13, alongside *AUTS2*, whereas canonical PRC1.2 core *PCGF2* and the PRC2 component *EED* declined (Figure S3C). These dynamics suggest that ncPRC1.3 components are specifically upregulated at the neural progenitor stage, leading us to hypothesize that FBRSL1-ncPRC1.3 directly engages corepressors at D13 to silence cell-cycle genes and enable neuronal differentiation.

### Loss of corepressor and ncPRC1.3 occupancy at neural progenitor promoters precedes cell-cycle de-repression

To define the chromatin basis of this developmental defect, we profiled CoREST, CTBP2, PCGF3, and RING1B genome-wide occupancy by ChIP-seq and chromatin accessibility by ATAC-seq in WT and p.Gln163* cells at D13. At the representative cell-cycle locus *CDCA8* (Cell Division Cycle Associated 8), WT neural progenitors displayed prominent promoter-proximal peaks of CoREST, CTBP2, PCGF3, and RING1B, with low chromatin accessibility (Figure 6A). In p.Gln163* progenitors, occupancy of CoREST, CTBP2, PCGF3, and RING1B was markedly lost, accompanied by a sharp gain in ATAC-seq promoter accessibility at D13 and subsequent de-repression of *CDCA8* mRNA at D17 (Figure 6A). Importantly, this loss of occupancy was selective, as robust binding was preserved at non-cell-cycle loci, including CTBP2 occupancy at *LHX2* and PCGF3 occupancy at *GATA6* (Figures S4A and S4B).

**Figure 6.**
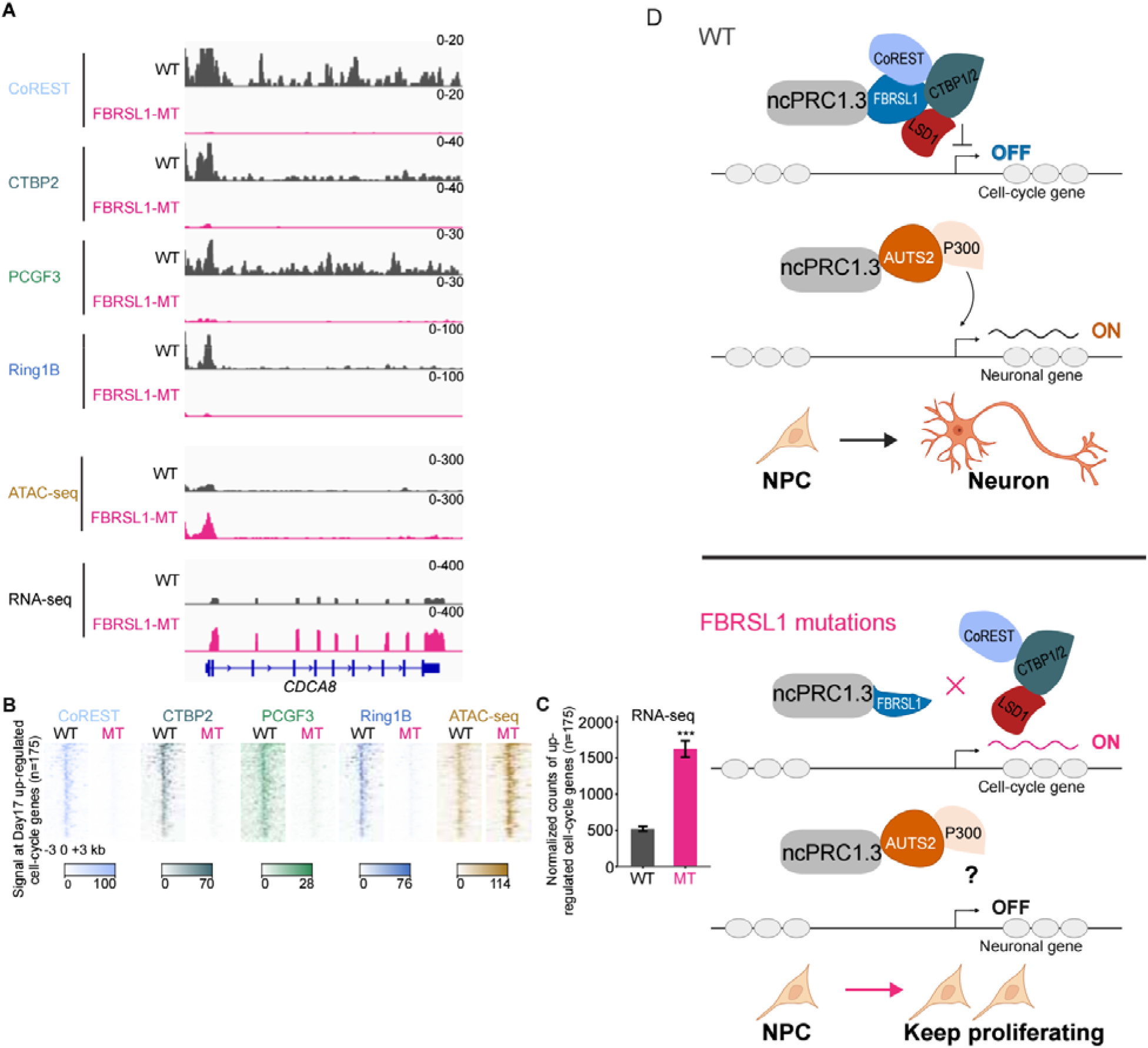
FBRSL1 p.Gln163* mutation disrupts CoREST/CtBP occupancy at cell-cycle genes, leading to their de-repression during the progenitor-to-neuron transition. **(A)** Genome browser view of the *CDCA8* locus. Tracks show ChIP-seq for CoREST, CTBP2, PCGF3 and RING1B, ATAC-seq at D13, and RNA-seq at D17, for WT and FBRSL1-MT. **(B)** Heat maps of ChIP-seq (CoREST, CTBP2, PCGF3 and RING1B) and ATAC-seq signal at D13 across ±3 kb of the transcription start site (TSS) of the 175 cell-cycle-related genes that are significantly up-regulated in FBRSL1-MT at D17. **(C)** D17 RNA-seq expression of the 175 cell-cycle genes in (B) in WT and FBRSL1-MT cells. Bars show mean ± SEM (n = 3). **(D)** Proposed model for FBRSL1-ncPRC1.3-mediated repression of cell-cycle genes through interaction with the CoREST/CTBP/LSD1 complex during the NPC-to-neuron transition. In WT cells at the NPC-to-neuron transition, FBRSL1–ncPRC1.3 recruits the CoREST/CTBP/LSD1 complex to repress cell-cycle genes while, based on prior work, AUTS2–ncPRC1.3 recruits p300 to activate neuronal genes, thereby facilitating neuronal differentiation. Patient-derived FBRSL1 truncations lack the Atrophin-1 domain and cannot engage the CoREST/CTBP/LSD1 complex, thereby maintaining cell-cycle genes in an active state. Through an unknown mechanism (?), neuronal genes also fail to be activated by AUTS2–ncPRC1.3, and progenitors keep proliferating, instead of becoming neurons.

To determine whether this chromatin defect is a genome-wide feature of the failure to silence proliferative programs, we examined the 175 cell-cycle-related genes that were significantly upregulated in mutant cells at D17. Heatmaps centered on the transcription start sites (TSS ±3 kb) of these 175 genes revealed intense promoter occupancy of CoREST, CTBP2, PCGF3, and RING1B in WT D13 progenitors (Figure 6B). In p.Gln163* progenitors, binding of all four factors collapsed to near-background levels, with a concomitant increase in promoter accessibility (Figure 6B). Analysis of the RNA-seq data confirmed that these 175 genes exhibited an average threefold elevated expression in mutant cells compared to WT at D17 (Figure 6C), documenting the magnitude of transcriptional de-repression following corepressor loss.

Collectively, these biochemical, genomic, and differentiation data establish an integrated model for ncPRC1.3 function in human corticogenesis (Figure 6D). In WT neural progenitors transitioning toward neuronal commitment, FBRSL1 couples ncPRC1.3 to the CoREST/CTBP/LSD1 corepressor complex to directly silence cell-cycle genes, allowing progenitors to exit proliferation. Once cell-cycle exit is licensed, the competing AUTS2-ncPRC1.3 subcomplex engages p300 to activate neuronal differentiation programs (Gao et al. 2014; Liu et al. 2021). In patient-derived p.Gln163* cells, the truncated FBRSL1 protein preserves PCGF3 binding but uncouples from corepressors, resulting in the failure of both corepressor and Polycomb machinery to engage with cell-cycle promoters. Consequently, cell-cycle genes fail to be repressed, and the neurogenic program fails to be executed, providing a molecular mechanism for the neurodevelopmental pathology observed in patients with *FBRSL1* mutations.

## Discussion

PcG complexes are canonically defined as transcriptional repressors, yet the noncanonical ncPRC1.3 assembly was paradoxically established as an activator through its incorporation of AUTS2 and resultant recruitment of p300 (Gao et al. 2014; Liu et al. 2021). Our findings resolve this paradox by demonstrating that ncPRC1.3 is an intrinsically versatile chromatin scaffold whose regulatory polarity is governed by mutually exclusive, paralogous adaptors. While AUTS2 equips the PCGF3 core with an activating arm, FBRSL1 binds the identical C-terminal RAWUL docking surface of PCGF3 to couple ncPRC1.3 to the CoREST/CTBP/LSD1 corepressor complex. This architecture explains how PCGF3-containing complexes can occupy both active and repressed genes and operate independently of classical Polycomb catalytic constraints (Scelfo et al. 2019). During human cortical differentiation, the failure of this repressive arm leaves proliferative gene programs sustained at the transition from neural progenitors to post-mitotic neurons, directly tying the molecular switch between paralogous adaptors to the execution of a key neurodevelopmental milestone.

The competitive displacement observed between AUTS2 and FBRSL1 for the PCGF3 RAWUL domain indicates that the stoichiometric balance between these paralogs dictates the transcriptional output of ncPRC1.3. In developing cortical tissue, this balance could be tuned by multiple physiological inputs. For instance, the deubiquitinase USP7 selectively represses *AUTS2* expression (Sijm et al. 2022), and neuronal activity dynamically modulates *Auts2* transcription (Pang et al. 2023). Precedents from other major chromatin remodelers demonstrate that paralog exchange is a widespread mechanism for developmental timing: subunit switching between BAF53a and BAF53b in BAF complexes, or exchange of SS18 for CREST, directly gates cell-cycle exit and neuronal differentiation in the developing cortex (Staahl et al. 2013; Braun et al. 2021), while developmental switching among CBX paralogs modulates canonical PRC1 targeting (O’Loghlen et al. 2012). In addition to AUTS2 and FBRSL1, the third paralog, FBRS, also co-purifies with the ncPRC1.3 core (Hauri et al. 2016; Gao et al. 2012). While FBRS exhibited neutral activity in our Gal4 reporter assay, a recent genome-wide screen established it as an essential component of ncPRC1.3 required for human naïve pluripotency reprogramming (Collier et al. 2022). Remarkably, PCGF3 preferentially partners with FBRS in naïve embryonic stem cells and switches to AUTS2 in primed cells (Collier et al. 2022). Thus, dynamic paralog switching on the PCGF3 scaffold appears to be an evolutionarily conserved mechanism utilized across diverse cell-fate transitions.

A central mechanistic question raised by our findings is how FBRSL1-ncPRC1.3 and its associated corepressors are targeted to specific genomic loci. In our human embryonic stem cell model, the patient-derived p.Gln163* truncation retains robust PCGF3 binding *in vitro*, yet neural progenitors carrying this mutation exhibit a concomitant loss of PCGF3, RING1B, CoREST, and CTBP2 occupancy at cell-cycle promoters. This demonstrates that docking to the ncPRC1.3 core is insufficient for stable promoter engagement *in vivo*; chromatin targeting strictly requires sequences downstream of codon 163 in FBRSL1. We envision two non-exclusive mechanisms for this localization. First, analogous to the recruitment of AUTS2-ncPRC1.3 by NRF1 (Liu et al. 2021), a sequence-specific transcription factor may interact with the central or C-terminal region of FBRSL1. A compelling candidate is the zinc-finger protein ZNF516, which was specifically recovered in our FBRSL1 interactome and is known to dock the CtBP/LSD1/CoREST complex to chromatin to repress proliferative genes (L. Li et al. 2017). Similarly, YY1 interacts with FBRSL1 (Kastens et al. 2025) and requires CtBP for Polycomb recruitment in *Drosophila* (Srinivasan and Atchison 2004), while atypical E2F family repressors, such as E2F7 repress cell-cycle promoters through direct CtBP recruitment. Second, corepressor engagement may provide essential avidity or cooperative stabilization: physical coupling to CoREST and CTBP2 could anchor ncPRC1.3 at target promoters, mirroring how ZNF198 stabilizes the LSD1-CoREST complex on chromatin (Gocke and Yu 2008), and REST stabilizes PRC1 at neuronal genes (Dietrich et al. 2012). Because endogenous FBRSL1 was refractory to direct ChIP-seq profiling, testing these targeting models represents an important objective for future study.

The molecular mechanism uncovered here provides a direct foundation for understanding *FBRSL1*-associated developmental pathology. All five reported pathogenic *FBRSL1* alleles converge upstream of the central Atrophin-1 domain (Ufartes et al. 2020; Pauli et al. 2021; Xu et al. 2026; Bukvic et al. 2024), and our biochemical analysis of p.Gln163* substantiates that this lesion constitutes a precise separation of function—severing corepressor recruitment while leaving the Polycomb scaffold intact. Although this corepressor uncoupling provides a molecular explanation for the neural progenitor cell-cycle defect and microcephaly, *FBRSL1* syndrome involves diverse systemic features that likely engage additional cellular pathways. Notably, FBRSL1 has been shown to interact with the spliceosome component SF3B1 to modulate *mdm2* mRNA splicing and p53-dependent apoptosis during cranial neural crest development, contributing to craniofacial dysmorphism (Gerstner et al. 2025). Furthermore, FBRSL1 associates with YY1 to support the expression of the chromatin regulators *BRPF1* and *KAT6A* in non-neural patient tissues (Kastens et al. 2025). Thus, FBRSL1 participates in distinct chromatin and post-transcriptional regulatory networks across embryonic development, with its corepressor-coupled ncPRC1.3 assembly functioning as the critical driver of human corticogenesis.

A striking feature of our differentiation model is that the failure to silence cell-cycle genes at Day 13 is followed by severely blunted induction of mature neuronal genes and an attenuated rise in *AUTS2* expression at Day 17 (Figure S3D). This phenotype highlights the temporal coordination between the two ncPRC1.3 arms. One primary explanation is that impaired *AUTS2* induction is a developmental consequence of failed neuronal commitment: AUTS2 is highly expressed in post-mitotic cortical neurons (Boucherie et al. 2025; Geng et al. 2024), and in the developing neocortex, the master neurogenic transcription factor TBR1 directly binds the *Auts2* promoter to activate its transcription (Bedogni et al. 2010). Because *TBR1* induction is virtually abolished in our mutant cells, the subsequent failure to induce *AUTS2* and its neurogenic program naturally follows. Alternatively, because AUTS2 actively promotes neuronal differentiation in human cerebral organoids and cortical progenitors (Boucherie et al. 2025; Fair et al. 2023; Geng et al. 2024), its diminished induction may actively compound the lineage stall. Notably, the logistics of these temporal events resolves a longstanding clinical paradox: mutations in *AUTS2* (an activator) and *FBRSL1* (a repressor) cause overlapping microcephaly and intellectual disability syndromes (Pauli et al. 2021). Our model indicates that these two paralogs operate in a sequential relay along a single developmental trajectory—FBRSL1-CoREST first extinguishes proliferative programs in neural progenitors, licensing AUTS2-p300 to subsequently activate neuronal differentiation programs. Disrupting either arm derails the identical developmental transition.

In conclusion, our study uncovers a two-state chromatin architecture in which the transcriptional output of ncPRC1.3 is dictated by opposing, paralogous adaptors. By coupling the Polycomb scaffold to the CoREST/CTBP/LSD1 corepressor complex, FBRSL1 provides a dedicated repressive mechanism required to silence proliferative programs and license human neuronal differentiation. Dissecting the precise upstream signals that govern the balance between FBRSL1 and AUTS2 will provide fundamental insights into the epigenetic control of human brain development and neurodevelopmental disease.

### Limitations of the study

Our conclusions are bounded by several technical and experimental limitations. First, our cellular model utilized homozygous p.Gln163* knock-in hESC lines to isolate the functional consequences of the truncated protein, whereas human patients carry heterozygous *de novo* mutations. While patient transcripts escape nonsense-mediated decay (Ufartes et al. 2020) and we confirmed stable mutant mRNA expression, technical limitations in detecting endogenous truncated FBRSL1 protein prevent us from distinguishing whether the clinical syndrome arises from pure haploinsufficiency or a dominant-negative mechanism. Second, direct genomic occupancy of endogenous FBRSL1 could not be mapped due to the lack of ChIP-grade antibodies, meaning that direct target loci are inferred from the localized loss of CoREST, CTBP2, PCGF3, and RING1B. Third, while genetic ablation demonstrated that CoREST and CTBP2 are strictly required for FBRSL1-mediated repression in reporter assays, the precise catalytic requirements for LSD1 demethylase or HDAC deacetylase activity were not tested, and corepressor requirements were not evaluated by acute endogenous depletion at native cell-cycle loci. Finally, paralog competition for the PCGF3 RAWUL domain was established using overexpression and co-immunoprecipitation; the absolute endogenous stoichiometry of AUTS2 versus FBRSL1 complexes across individual differentiating cells remains to be determined.

## STAR Methods

### KEY RESOURCES TABLE

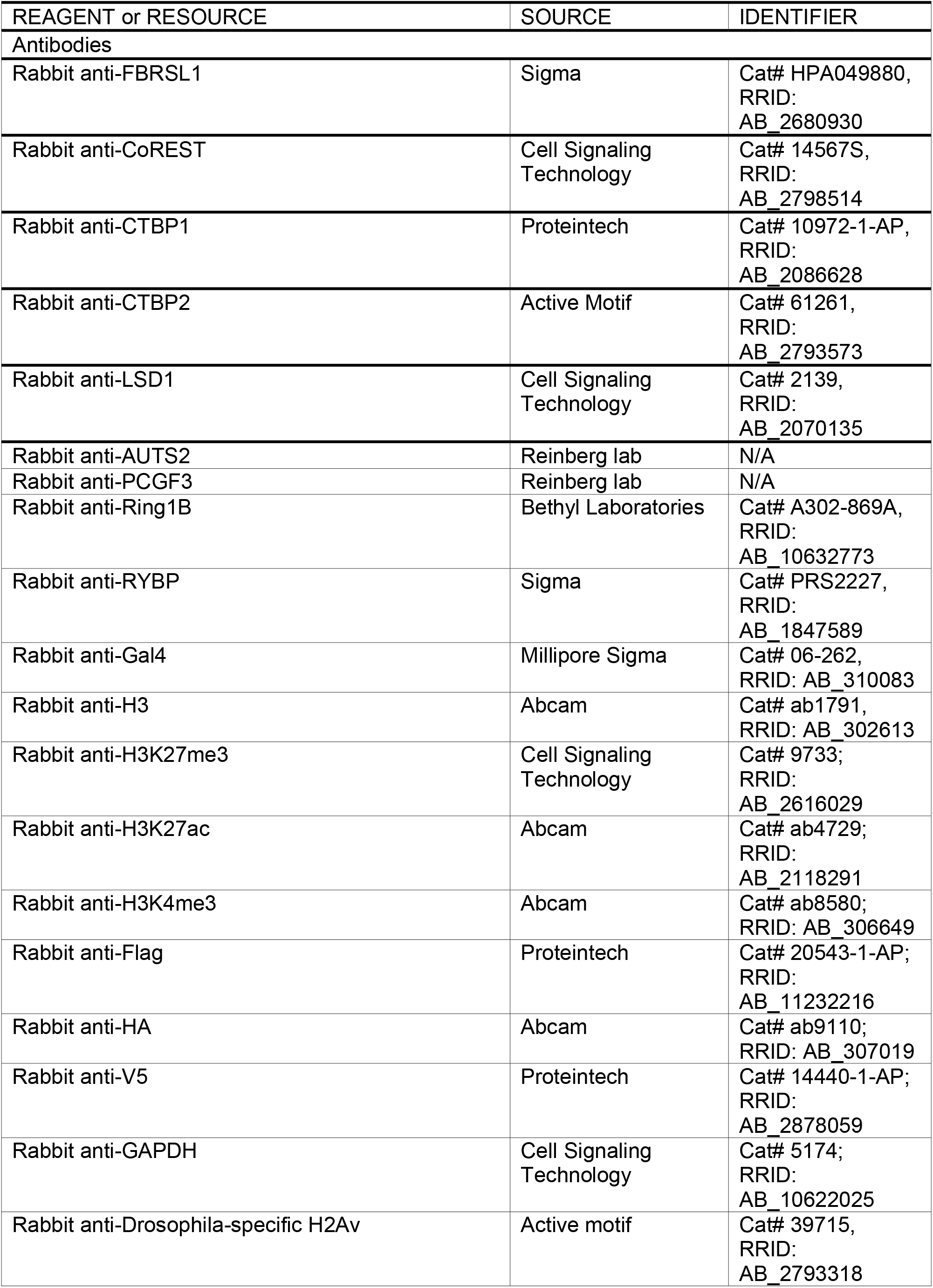
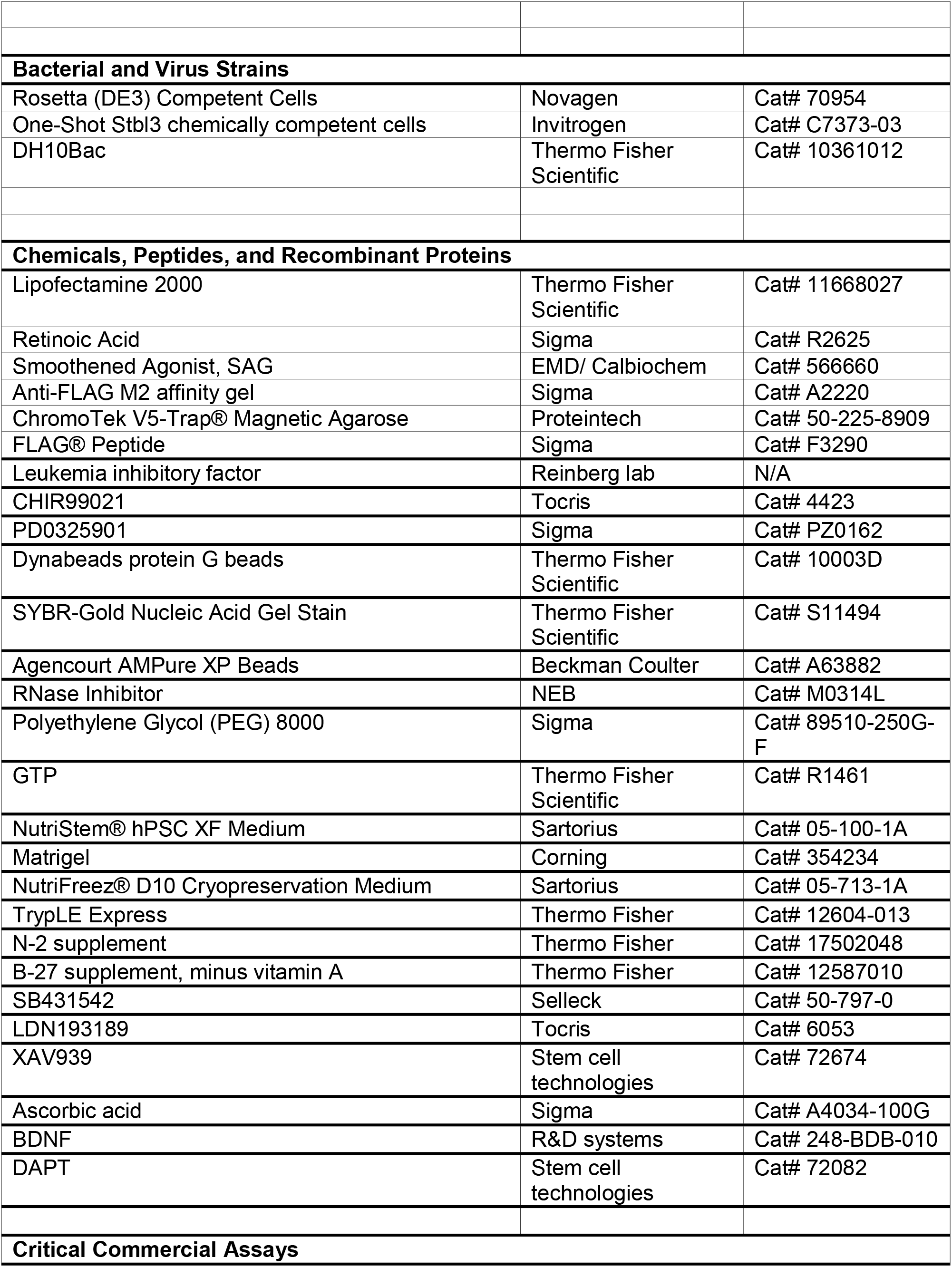
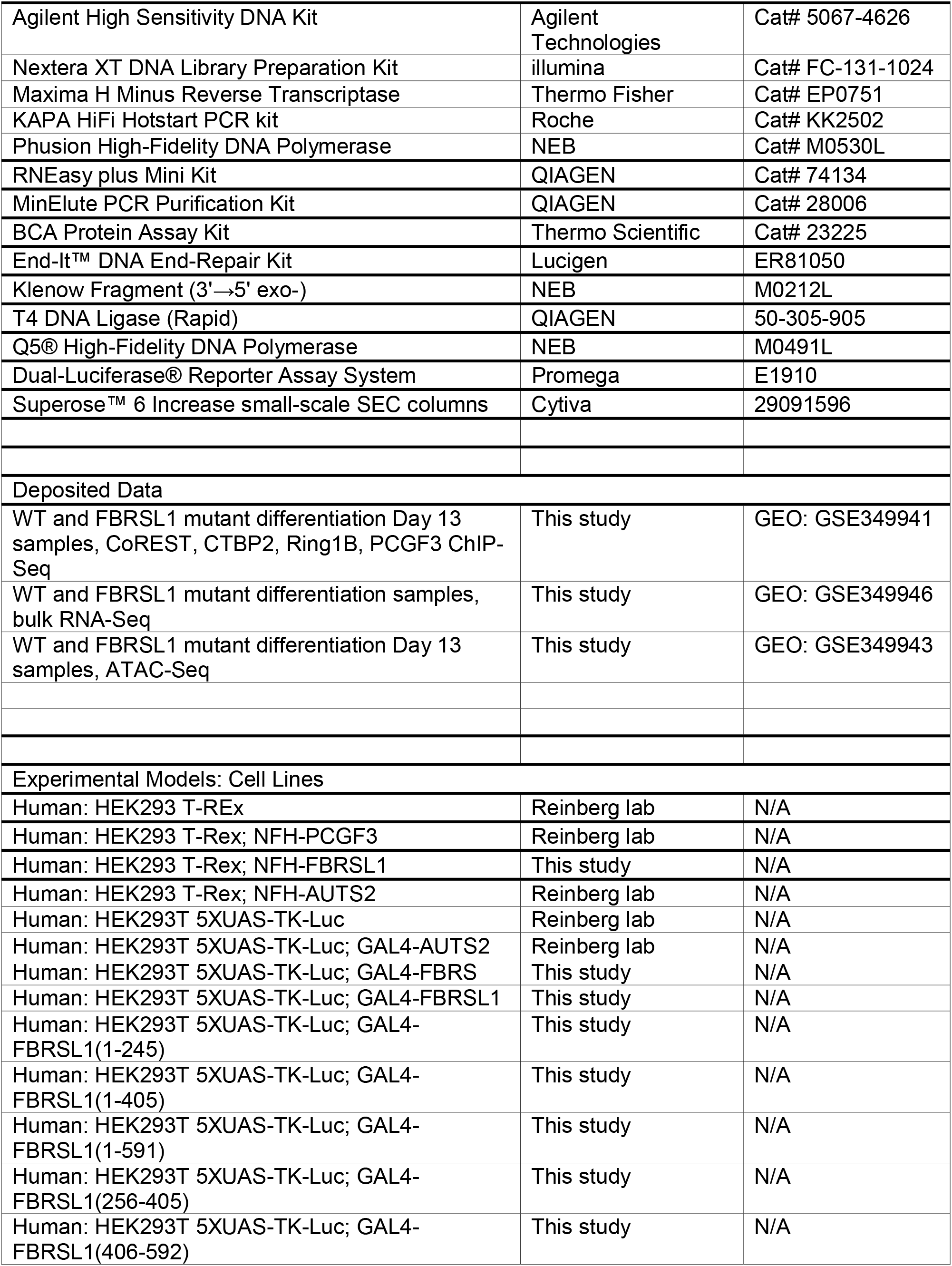
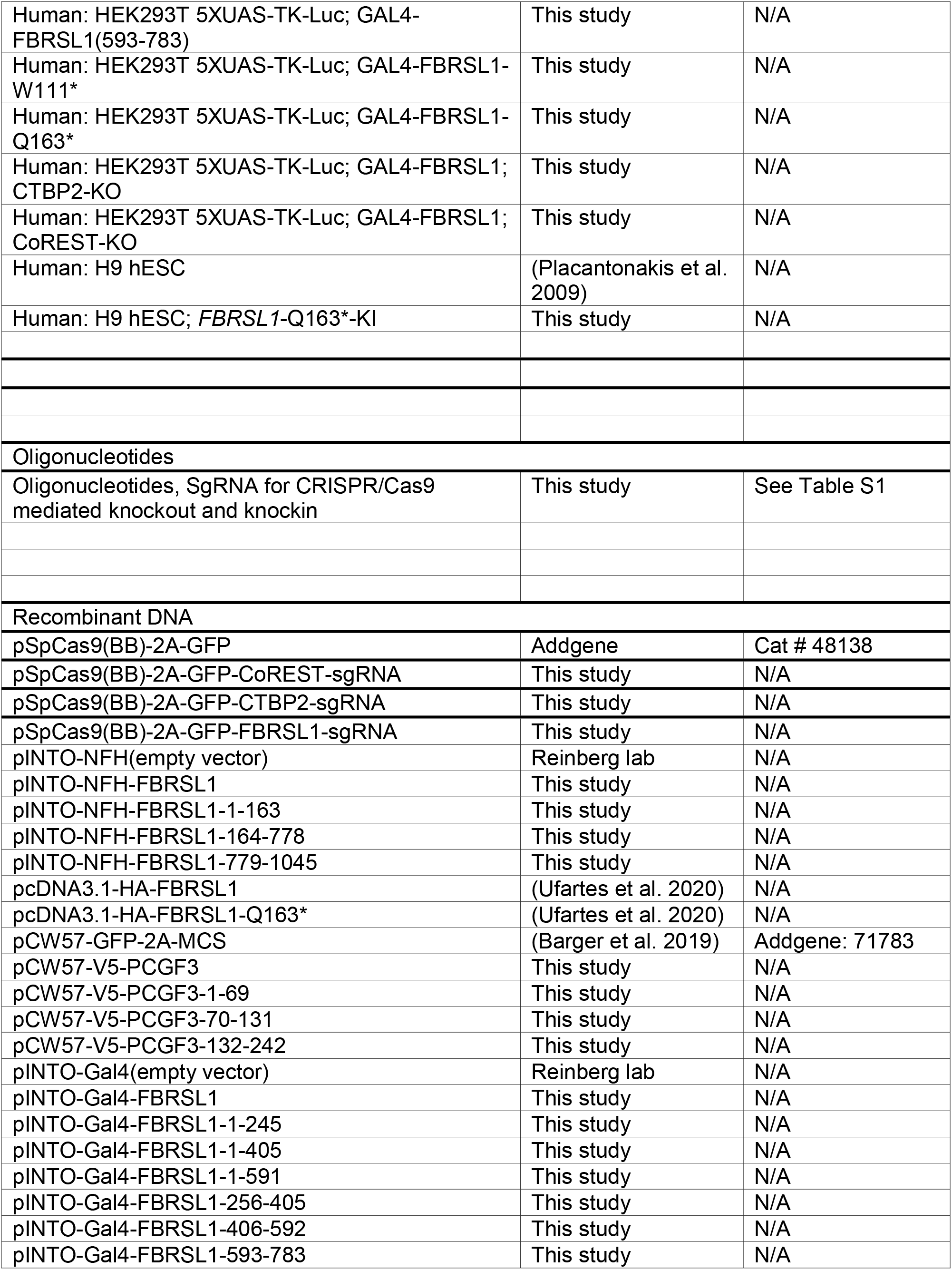
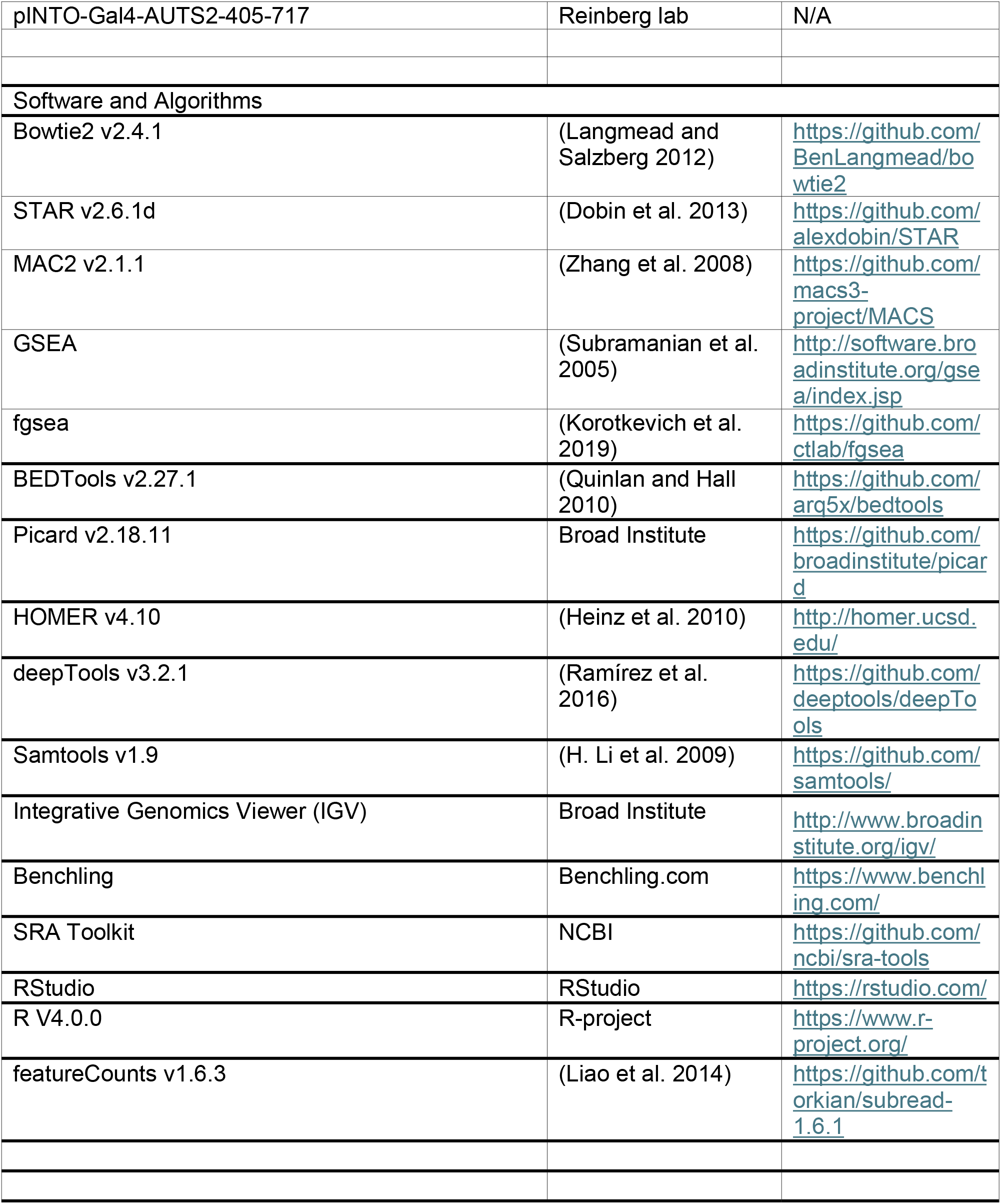

### Lead Contact and Materials Availability

Please direct inquiry to the Lead Contact, Danny Reinberg. All unique/stable reagents generated in this study are available from the Lead Contact with a complete Materials Transfer Agreement.

### Experimental Model and Subject Details

#### Cell lines and culture condition

All hESC lines (H9 and derivatives) were routinely cultured feeder-free on culture vessels pre-coated with growth factor-reduced Matrigel (Corning; diluted 1:100 in DMEM) at 37°C in a humidified incubator with 5% CO. Cells were maintained in NutriStem® hPSC XF Medium (Sartorius / Biological Industries) with daily medium replacement. hESCs were passaged upon reaching ∼75%–80% confluency using 0.5 mM EDTA in sterile PBS (pH 7.4). Cells were gently dissociated and replaced as small multicellular aggregates without enzymatic digestion to maintain genomic stability and pluripotency.

HEK293 T-Rex and HEK293 T-Rex 5XGal4TK-Luc cells were cultured in standard DMEM supplemented with 10% FBS, 100 U/mL penicillin-streptomycin.

### Method Details

#### Immunoprecipitation, fractionation and Proteomics

Cells were lysed on ice for 5 min in hypotonic HMSD buffer (20 mM HEPES, pH 7.5, 5 mM MgCl, 250 mM sucrose, 25 mM NaCl, 1 mM DTT) supplemented with protease inhibitors (0.2 mM PMSF, 1 µg/mL each of pepstatin A, leupeptin, and aprotinin) and phosphatase inhibitors (10 mM NaF and 1 mM Na VO). Nuclei were pelleted at 1,000 × g for 5 min at 4°C, washed once in HMSD buffer, and resuspended in high-salt BC420 buffer (20 mM Tris-HCl, pH 7.9, 0.42 M NaCl, 1.5 mM MgCl, 0.2 mM EDTA, 0.5 mM DTT, with protease and phosphatase inhibitors). Nuclear lysis proceeded at 4°C for 1 h. After clearing insoluble debris at 20,000 × g for 15 min at 4°C, supernatants were dialyzed overnight at 4°C against Buffer D (20 mM HEPES, pH 7.9, 150 mM NaCl, 1.5 mM MgCl, 0.2 mM EDTA, 5% glycerol, 1 mM DTT, and 0.2 mM PMSF). Extracts were clarified at 20,000 × g for 10 min, and protein concentrations were quantified via bicinchonic acid (BCA) assay.

To isolate native FBRSL1 complexes, dialyzed nuclear extracts were applied to a Phosphocellulose P11 (Whatman) column equilibrated in Buffer BC100 (20 mM Tris-HCl, pH 7.9, 100 mM NaCl, 1.5 mM MgCl, 0.2 mM EDTA, 10% glycerol, 1 mM DTT, and 0.2 mM PMSF). Bound proteins were resolved via stepwise elution using Buffer BC containing 0.1 M, 0.3 M, 0.5 M, and 1.0 M NaCl. The 0.5 M NaCl eluate containing FBRSL1 was adjusted to ∼150 mM NaCl by dilution with salt-free Buffer BC0 and incubated with anti-FLAG M2 affinity resin (Sigma-Aldrich) overnight at 4°C. After extensive washing with Buffer D, bound complexes were competitively eluted under native conditions with 0.5 mg/mL 3× FLAG peptide (Sigma-Aldrich) in Buffer D for 1 h at 4°C. For native molecular mass determination, 3× FLAG peptide eluates were concentrated using an Amicon Ultra centrifugal filter (10-kDa MWCO; Millipore) and fractionated on a Superose 6 column (Cytiva) equilibrated with Buffer D on an ÄKTA Pure FPLC system.

Fractions were collected across the total column volume and analyzed by SDS-PAGE and western blotting.

For analytical endogenous co-immunoprecipitation, 1–2 mg of clarified nuclear extract was incubated with 1–3 µg of specific primary antibody or control IgG in Buffer D for 2 h at 4°C, followed by overnight capture with 30 µL of Protein G agarose beads. Beads were washed three times in Buffer D, and bound proteins were eluted with 0.2 M glycine-HCl (pH 2.6; immediately neutralized with 1 M Tris-HCl, pH 8.5) or denatured in 1× SDS sample buffer at 95°C.

For proteomic profiling, affinity-purified complexes or immunoprecipitates were resolved on 4%– 12% NuPAGE Bis-Tris gels (Invitrogen) and visualized by Coomassie Brilliant Blue. Gel bands (or eluates) were subjected to in-gel reduction (10 mM DTT, 56°C), alkylation (55 mM iodoacetamide, RT, in the dark), and digested overnight at 37°C with sequencing-grade modified trypsin (Promega), followed by standard LC/MS/MS procedure.

#### Whole cell extract and western blotting

Cells were harvested and lysed with CHAPS-Urea buffer (50 mM Tris-HCl, pH 7.9, 8M Urea, and 1% CHAPS) containing protease inhibitors and phosphatase inhibitors as mentioned above. The cell suspension was briefly sonicated (40% amplitude, 5 strokes) and centrifuged at 20,000 x g at 4°C for 20 min. The supernatant was collected and protein concentrations were quantified via BCA assay. Proteins were separated using a 6%–12% SDS PAGE gel and transferred onto a PVDF membrane. Membranes were blocked with 5% milk in PBST at RT for 1 h and incubated with primary antibody overnight at 4°C. Membranes were washed 3 times with PBST and then incubated with HRP-conjugated secondary antibodies for 1 h at RT, followed by exposure.

#### CRISPR-mediated genome editing

Single-guide RNAs (sgRNAs) targeting CoREST (*RCOR1*), *CTBP2*, and *FBRSL1* were designed using the Benchling CRISPR design platform to minimize predicted off-target cleavage (Table S1). All sgRNAs were cloned into the BbsI restriction site of the vector pSpCas9(BB)-2A-GFP (PX458; Addgene #48138, gift from Feng Zhang).

To generate knockout (KO) models, HEK293 T-REx cells at ∼70% confluency were transfected with sequence-verified PX458-sgRNA targeting plasmids using linear polyethylenimine (PEI; 1:3 DNA:PEI ratio). At 48 h post-transfection, single GFP-positive cells were isolated by fluorescence-activated cell sorting (FACS), and 20,000 cells were plated on a 15 cm dish. Single cells were allowed to grow to a colony for ∼7 days and then picked, trypsinized in Accutase for 5 min, and split into two individual wells of two 96-well plates for genotyping and culture respectively. Genomic DNA was extracted in lysis buffer (50 mM Tris-HCl, pH 8.0, 2 mM NaCl, 10 mM EDTA, and 0.1% SDS) supplemented with 200 µg/mL Proteinase K at 55°C for 2 h, followed by heat inactivation at 95°C. Targeted loci were PCR-amplified using flanking primers (Table S1) and verified for frameshift indel mutations by Sanger sequencing and ICE/TIDE deconvolution analysis. Complete loss of target protein expression was validated by western blotting.

The patient-derived *FBRSL1* pathogenic variant was engineered into the endogenous locus of H9 hESCs via homology-directed repair (HDR). The HDR donor template was synthesized as an antisense single-stranded oligodeoxynucleotide (ssODN) featuring 5′ biotinylation to enhance knock-in fidelity, centered on the point mutation and silent blocking mutations within the PAM or seed sequence to prevent re-cleavage. Cultured H9 hESCs (∼80% confluency) were pre-conditioned with 2 µM ROCK inhibitor (Y-27632) for 2–4 h, dissociated into single cells using TrypLE for 6 min at 37°C, and passed through a 100-µm cell strainer. Electroporation was performed using the Neon™ Transfection System (Invitrogen). Briefly, 5M dissociated hESCs were collected by centrifugation (200 × *g*, 3 min, 4°C), resuspended in 100 µL of Buffer R, and mixed with 3 µg of *FBRSL1*-targeting PX458 plasmid and the 5′-biotin-modified ssODN template. Cells were electroporated in a 100-µL Neon Pipette Tip applying a single pulse at 1,350 V with a 30-ms pulse width. Following electroporation, cells were immediately seeded onto pre-warmed, Matrigel-coated plates containing recovery medium. Complete medium without ROCK inhibitor was replenished 24 h post-electroporation. At 48 h post-nucleofection, cells were sorted, planted at a low density, and single colonies were picked, expanded and validated as described above.

#### hESC to Cortical Neuron Differentiation

Cortical neural differentiation was adapted from established dual-SMAD and canonical Wnt inhibition strategies. Briefly, two days prior to induction (Day −2), H9 hESCs grown to ∼85% confluency in a 10-cm dish were washed once with PBS and dissociated into small aggregates using 5 mL PBS containing 0.5mM EDTA for 5 min at RT. EDTA was removed and cells were collected and re-seeded at high density into Matrigel-coated 6-well plates at a 1:3 split ratio (3 mL per well of NutriStem containing 10 µM Y-27632). On Day −1, medium was replaced with 3 mL of fresh NutriStem without ROCK inhibitor.

Neural induction was initiated on Day 0. Cells were washed twice with PBS and switched to 3 mL of basal N2B27 induction medium consisting of a 1:1 mixture of: DMEM/F-12 with GlutaMAX™ containing 1× N2 supplement (final dilution 1:200), Neurobasal™ Medium containing 1× B-27™ supplement minus vitamin A (final dilution 1:100), and 0.1 mM β-mercaptoethanol (1:1,000).

On Days 0, 2, and 4, this basal N2B27 medium was freshly supplemented with small molecules targeting neural lineage commitment: 10 µM SB431542, 1 µM LDN-193189, and 3 µM XAV939. Prior to feeding on Days 2 and 4, wells were washed twice with PBS. On Day 6, cultures were washed twice with PBS and replenished with 3 mL of basal N2B27 medium supplemented with 10 µM SB431542 and a reduced concentration of LDN-193189 (250 nM), with complete withdrawal of XAV939. On Day 8, 3 mL of fresh basal N2B27 medium was directly added to each well without discarding the spent medium, yielding conditioned medium for subsequent dissociation and replating.

On Day 9, the conditioned medium was harvested, supplemented with 10 µM ROCK inhibitor (Y-27632), and filtered. Adherent human neural stem cells (hNSCs) were washed once with PBS and incubated with 2 mL of TrypLE™ Express per well for 3–4 min at 37°C. Cells were gently detached and collected into 2 mL of pre-warmed DMEM/F-12 (GlutaMAX™), followed by centrifugation at 1,000 rpm for 3 min. The resulting cell pellet was resuspended in 1.5 mL of the collected conditioned medium. Viable cells were quantified and adjusted to a final density of 1×10^7^ cells/mL in conditioned medium containing 10 µM Y-27632.

Fresh 6-well plates were pre-coated with Matrigel (1:100 in DMEM) for 1 h at 37°C. After aspirating the Matrigel solution, a circular perimeter was demarcated in the center of each well using a sterile pipette tip, and a 400-µL droplet (about 4M cells) was carefully deposited within the delineated zone to establish a high-density monolayer. Plates were placed in a 37°C incubator for 15 min to promote rapid cellular adherence.

During cell attachment, neuronal maturation medium was prepared by diluting the remaining conditioned medium 1:3 with fresh basal N2B27 medium (1-part conditioned medium to 2 parts fresh N2B27), supplemented with: 0.2 mM L-ascorbic acid, 10 ng/mL recombinant human BDNF, and 10 µM DAPT. After the 15-min attachment, 2 mL of maturation medium was gently introduced along the side wall of each well to avoid dislodging the high-density cell patch.

On Days 11 and 13, wells were washed twice with PBS and replenished with 2 mL of fresh basal N2B27 medium supplemented with 0.2 mM L-ascorbic acid, 10 ng/mL BDNF, and 10 µM DAPT. Distinct, morphologically mature cortical neurons with extensive neurite arborization emerged between Days 14 and 15, and cultures were maintained through Day 20 for downstream experimental assays.

#### Luciferase reporter assay

All human *FBRSL1* constructs used in this study were generated based on the cDNA sequence matching NCBI Reference Sequence NM_001142641.2. HEK293T 5XGal4 TK-Luc cells stably transfected with pINTO-GAL4 vector control or with inserts of interest were treated with 100 ng/ml doxycycline. Cells were lysed by adding 250 ul of ice-cold lysis buffer (0.2% Triton X-100, 100mM potassium phosphate, pH 7.8, and 1mM DTT) and shaking for 10 min at 4°C. The cell lysate was centrifuged at 20,000g for 10 min and the protein concentration of the resulting supernatant was determined by BCA assay. 30 ug of the supernatant was assayed for luciferase activity using luciferase assay substrate (Promega).

#### ChIP-qPCR and ChIP-seq library preparation

For chromatin preparation, HEK293 T-REx cells were detached using 0.05% trypsin-EDTA, whereas H9-derived cortical lineage cultures were collected at defined differentiation stages (Days 0, 4, 9, 13, and 17) using TrypLE™ Express. Approximately 20M dissociated cells were subjected to dual cross-linking: cells were first resuspended in 10 mL PBS containing 2 mM disuccinimidyl glutarate (DSG; diluted from freshly made 0.5 M solution) and incubated for 30 min at RT with gentle rotation. Cells were pelleted at 500 × g for 4 min, washed twice with PBS, and subsequently cross-linked in 10 mL of 1% paraformaldehyde (PFA) in PBS for 10 min at RT. Fixation was quenched with 125 mM glycine for 5 min, followed by two washes in ice-cold PBS.

Nuclei were isolated using buffers in the following order: LB1 (50 mM HEPES, pH 7.5 at 4°C, 140 mM NaCl, 1 mM EDTA, 10% Glycerol, 0.5% NP40, and 0.25% Triton X; 10 min at 4°C), LB2 (10 mM Tris, pH 8 at 4°C, 200 mM NaCl, 1 mM EDTA, and 0.5 mM EGTA; 10 min at 4°C), and LB3 (10 mM Tris, pH 7.5 at 4°C, 1 mM EDTA, 0.5 mM EGTA, and 0.5% N-Lauroylsarcosine sodium salt). Chromatin was fragmented to an average size of 250 bp in LB3 buffer using a Diagenode Bioruptor. 200 μg sonicated chromatin, 4 ug antibody and 20ul Dynabeads were used in each ChIP reaction supplemented with 0.5x volume of incubation buffer (3% Triton X, 0.3% Na Deoxycholate, and 15 mM EDTA). 1 μg of Drosophila chromatin and 0.2 μg of anti-Drosophila H2A.X antibody were added in each ChIP reaction as spike-in references. After washing 5X with RIPA buffer (50 mM HEPES, pH 7.5 at 4°C, 0.7% Na Deoxycholate, 1 mM EDTA, 1% NP40, and 500 mM LiCl) and once with TE+50 mM NaCl, beads-bound DNA was eluted in freshly prepared elution buffer (50 mM Tris, pH 8, 10 mM EDTA, and 1% SDS) at 65°C for 20 min. Eluted DNA was de-crosslinked at 65°C overnight, followed by protease K and RNase A treatment.

For ChIP-qPCR analysis in HEK293 T-REx cells, purified ChIP DNA and a 1:100 dilution of input DNA (1% equivalent) was amplified in triplicate using SYBR Green qPCR master mix with primers specifically targeting the upstream activation sequence (UAS) element.

For Library preparation, IP’ed DNA (∼1-30 ng) was end-repaired using End-It Repair Kit, tailed with deoxyadenine using Klenow exo-, and ligated to custom adapters with T4 Rapid DNA Ligase (Enzymatics). Fragments of 200-600 bp were size selected using Agencourt AMPure XP beads (0.5X and 0.3X) and subjected to PCR amplification using Q5 DNA polymerase. Libraries were size selected using AMPure XP beads (0.75X), quantified by Qubit™ dsDNA HS Assay Kit and quality checked by High Sensitivity D1000 ScreenTape. Libraries were sequenced as 150 bp paired end reads on the Illumina NovaSeq X platform.

#### RNA-seq library preparation

Transcriptome profiling was performed using the bulk-adapted FLASH-seq protocol (Hahaut et al. 2022; Siwicki et al. 2023, v. 2), which enables sensitive, full-length cDNA generation and library construction from low-input RNA samples.

Total RNA was extracted from harvested cells using the RNeasy Plus Mini Kit (QIAGEN) with on-column genomic DNA elimination. Purified RNA was normalized to 1–2 ng/µL in nuclease-free water. RNA aliquots (1) were combined with a lysis and annealing mixture containing 0.2% Triton X-100, recombinant RNase inhibitor (Takara Bio), 2.5 mM dNTPs, and 2.5 µM anchored oligo(dT) primer harboring an ISPCR universal handle. Secondary RNA structures were denatured, and primer annealing was carried out by incubating at 72°C for 3 min, followed by immediate cooling on ice.

Reverse transcription and cDNA pre-amplification were performed in a coupled reaction. The annealed RNA was supplemented with a master mix consisting of SuperScript™ IV Reverse Transcriptase (200 U/µL; Thermo Fisher Scientific), 2 µM template-switching oligonucleotides (TSO; containing 3’ locked nucleic acid [LNA] residues), and 2x KAPA HiFi HotStart ReadyMix (Roche). Reverse transcription was conducted at 50°C for 60 min, followed by enzyme inactivation at 98°C for 3 min. Full-length cDNA was immediately amplified by 14–16 cycles of PCR (98°C for 20 s, 67°C for 20 s, and 72°C for 6 min) using ISPCR primers, terminating with a final extension at 72°C for 5 min.

Amplified double-stranded cDNA was purified using a 0.8× volumetric ratio of SPRI magnetic beads (AMPure XP, Beckman Coulter) and eluted in nuclease-free water. cDNA yield and size distribution (peaking at ∼1.5–2.0 kb) were determined using the Qubit™ dsDNA HS Assay Kit (Invitrogen) and High Sensitivity D5000 ScreenTape on an Agilent 4200 TapeStation.

For sequencing library construction, 1 ng of purified full-length cDNA (normalized to 200 pg/µL) was tagmented using home-made Tn5 transposase loaded with adapter sequences for 10 min at 55°C. Tagmentation was terminated by the addition of 0.2% SDS. Dual-indexed sequencing adapters were incorporated via indexing PCR using 1 µM unique dual-indexed Nextera primers and KAPA HiFi HotStart ReadyMix (12 cycles). The resulting libraries were purified using 0.8× AMPure XP beads to eliminate primer dimers and residual fragments <200 bp. Library molarity and fragment distribution (average size ∼350-450 bp) were quantified using Qubit dsDNA HS and High Sensitivity D1000 ScreenTape. Indexed libraries were pooled equimolarly and sequenced to a target depth of ≥25 million reads per sample in 150-bp paired-end mode on an Illumina NovaSeq X platform.

#### ATAC-seq library preparation

Chromatin accessibility profiling was conducted using the Omni-ATAC protocol (Corces et al. 2017). For each sample, 50,000 viable cells were harvested and pelleted at 500 × *g* for 5 min at 4°C in a fixed-angle centrifuge. The supernatant was carefully aspirated, and the cell pellet was gently resuspended in 50 µL of ice-cold ATAC-Resuspension Buffer (ATAC-RSB: 10 mM Tris-HCl, pH 7.4, 10 mM NaCl, and 3 mM MgCl) supplemented with 0.1% NP-40, 0.1% Tween-20, and 0.01% digitonin. Cells were lysed on ice for 3 min. Lysis was immediately stopped by adding 1 mL of ice-cold ATAC-RSB containing 0.1% Tween-20 (without NP-40 or digitonin), followed by gentle tube inversion. Intact nuclei were pelleted at 500 × *g* for 10 min at 4°C, and the supernatant was aspirated.

Isolated nuclei were resuspended in 50 µL of transposition reaction mixture consisting of: 25 µL 2× TD Buffer (20 mM Tris-HCl, pH 7.6, 10 mM MgCl, and 20% [v/v] dimethylformamide [DMF]), 2.5 µL Tn5 transposase (100 nM final concentration; home-made), 16.5 µL PBS, 0.5 µL 1% digitonin (0.01% final), 0.5 µL 10% Tween-20 (0.1% final), and 5 µL nuclease-free water. Transposition was carried out at 37°C for 30 min in a thermomixer with continuous agitation at 1,000 rpm. Transposed DNA fragments were immediately purified using the Zymo DNA Clean & Concentrator-5 Kit (Zymo Research) and eluted in 21 µL of elution buffer.

Transposed DNA (20 µL) was amplified for 5 cycles in a 50-µL reaction using NEBNext® High-Fidelity 2× PCR Master Mix (NEB) and 1.25 µM customized indexed primers (Ad1 and barcoded Ad2) under the following conditions: 72°C for 5 min, 98°C for 30 s, followed by 5 cycles of 98°C for 10 s, 63°C for 30 s, and 72°C for 1 min.

Amplified libraries were purified using the Zymo DNA Clean & Concentrator-5 Kit and eluted in 20 µL of nuclease-free water. Final library concentrations and library fragment size were evaluated on an Agilent 4200 TapeStation using High Sensitivity D1000 ScreenTape. Equimolar library pools were sequenced to generate 150-bp paired-end reads on an Illumina NovaSeq X platform.

### Quantification and Statistical Analysis

#### RNA-seq data analysis

Reads were aligned to the human reference genome (GRCh38/hg38) with STAR (v2.7.11b) in two-pass mode, using a GENCODE v49 splice-junction database and the parameters ‘-- outFilterMismatchNoverLmax 0.2 --outFilterMultimapNmax 1 --outFilterType BySJout -- outSAMstrandField intronMotif --outSAMmapqUnique 60 --twopassMode Basic -- outSJfilterReads Unique --outFilterIntronMotifs RemoveNoncanonical’, so that only uniquely mapped reads were retained. Fragments were assigned to genes with featureCounts (Subread; ‘-p -s 0 -t exon’) against the GENCODE v49 annotation.

Analyses used WT and FBRSL1-MT libraries at five time points, with three biological replicates per genotype and time point (30 libraries). Genes with fewer than 10 counts in at least 3 libraries were removed. Count matrices were analysed in DESeq2 using a single-factor design (’∼ group’, group = genotype × time point). Size-factor normalization and dispersion estimation were performed by DESeq2, and Wald tests were run for FBRSL1-MT versus WT at each time point. P values were adjusted by the Benjamini–Hochberg method, and log2 fold changes were shrunk with the ashr estimator. Differentially expressed genes (DEGs) were defined as adjusted P < 0.05 and |log2 fold change| ≥ 1. Variance-stabilized (VST) counts were used for PCA and for visualization of module-level trajectories.

Trajectories of lineage-marker genes (pluripotency, neural progenitor and cortical neuron markers) and of selected Polycomb/PRC1.3 complex genes were plotted as DESeq2 size-factor-normalized counts across the five time points (individual replicates and group means). For module-level trajectories, 12 curated genes per module (cell cycle, cortical neuron function) were z-scored across all libraries on VST values, averaged per library, and plotted as mean ± s.e.m. across the three replicates.

Gene set enrichment analysis was performed with fgsea on all tested genes ranked by the DESeq2 Wald statistic (unshrunk), separately for each time point (FBRSL1-MT versus WT), using MSigDB (msigdbr, Homo sapiens) Gene Ontology Biological Process (C5:GO:BP) gene sets.

The cell-cycle gene set was defined as genes significantly up-regulated in FBRSL1-MT relative to WT at Day 17 (adjusted P < 0.05, log2 fold change ≥ 1) that were also members of the MSigDB gene sets HALLMARK_E2F_TARGETS, HALLMARK_G2M_CHECKPOINT, REACTOME_CELL_CYCLE_MITOTIC or REACTOME_CELL_CYCLE_CHECKPOINTS (n = 175 genes). The neuronal-function gene set comprised significantly down-regulated Day 17 DEGs belonging to REACTOME_NEURONAL_SYSTEM, GOBP_REGULATION_OF_NEURON_PROJECTION_DEVELOPMENT or GOBP_CENTRAL_NERVOUS_SYSTEM_NEURON_DIFFERENTIATION. For the 175 cell-cycle genes, DESeq2-normalized expression in WT and mutant cells at Day 17 was compared by two-sided Wilcoxon rank-sum test.

#### ChIP-seq data analysis

Reads were trimmed with fastp. Each trimmed read pair was aligned independently to the human genome (hg38) and to the D. melanogaster genome (dm6) with Bowtie2 (v2.3.5.1), using separately optimized alignment modes for the two genomes:

- hg38: ‘--local --very-sensitive --no-mixed --no-discordant -I 10 -X 700’.
- dm6 (spike-in): ‘--end-to-end --very-sensitive --no-overlap --no-dovetail --no-mixed --no-discordant -I 10 -X 700’.

Alignments with MAPQ < 20 were removed with samtools (v1.9), PCR duplicates were identified with Picard MarkDuplicates (v2.18.11) and reads overlapping the ENCODE hg38 blacklist were removed with bedtools (v2.30.0). Duplicate-flagged reads were excluded from all downstream signal tracks and peak calling.

Signal tracks (bigWig, 50 bp bins) were generated with deepTools (’bamCoverage --binSize 50 - -normalizeUsing None --ignoreForNormalization chrX’) and scaled by a per-library spike-in scale factor derived from the number of uniquely aligned, de-duplicated dm6 fragments, following the principle of exogenous-reference normalization (Orlando et al. 2014). Peaks were called with MACS3 (’callpeak -f BAMPE -g hs --keep-dup all -q 0.05’) against the WT input, in narrow-peak mode for transcription-factor and chromatin-regulator ChIPs.

Promoters were defined as TSS ± 2 kb (GENCODE v49, clipped to chromosome bounds) and mean spike-in-normalized signal per promoter was extracted with deepTools multiBigwigSummary. For heatmaps, signal was computed with computeMatrix (reference-point at the TSS, ±3 kb, 50 bp bins) and plotted with plotHeatmap for 175 Day-17 cell-cycle up-regulated genes. Genes were ordered by descending WT CTBP2 signal across the same window in all panels.

#### ATAC-seq data analysis

Reads were trimmed with fastp. Trimmed reads were aligned to hg38 with Bowtie2 (v2.3.5.1; ‘-- local --very-sensitive --no-mixed --no-discordant -I 10 -X 2000’). Alignments with MAPQ < 30 were removed (samtools v1.9), PCR duplicates were removed with Picard MarkDuplicates (’REMOVE_DUPLICATES=truè), and reads mapped to the mitochondrial genome or within ENCODE hg38 blacklist regions were excluded. Read positions were adjusted for Tn5 insertion offsets (+4 bp on the plus strand, −5 bp on the minus strand) using deepTools alignmentSieve ‘- -ATACshift’. Fragment-size distributions and the fraction of reads in peaks (FRiP) were used as quality metrics.

Accessible regions were called per library with MACS3 (’callpeak -f BAMPE -g hs --keep-dup all -q 0.05’). Tracks were generated with deepTools bamCoverage (’--binSize 10 –normalizeUsing RPKM --ignoreForNormalization chrX’); replicate tracks were averaged per genotype with bigwigCompare (’--operation mean’, 50 bp bins).

Promoter accessibility was quantified as the mean RPKM signal within TSS ± 2 kb for each library. Heatmaps of Day 13 ATAC signal at the 175 Day-17 cell-cycle up-regulated genes were generated with computeMatrix (reference-point at the TSS, ±3 kb, 50 bp bins) and plotHeatmap.

## Data and Code Availability

All sequencing datasets generated as part of this study will be publicly available in NCBI GEO and are available for the review process with the provided token access key (GSE349941 (token: yxwfyyuwpvqdnmz), GSE349943 (token: stgbuoyodnqtlml), GSE349946 (token: sjivgaqyzdypjct)). Further information and requests for resources and reagents should be directed to the lead contact, Danny Reinberg. The publicly available resources used (reference genome sequence and annotation, and computational analysis tools) are described in the Methods section. Only publicly available tools were used in data analysis. The analysis parameters used have been described in Methods.

## Acknowledgments

We thank Dr. Lynne Vales for critical editing and insightful review of the manuscript, and past and current members of the Reinberg laboratory for helpful discussions and constructive suggestions. We are grateful to Dr. Silke Pauli and members of the Pauli laboratory (University Medical Center Göttingen, Germany) for valuable discussions regarding patient-derived *FBRSL1* variants and for generously sharing mutant plasmid expression vectors. We thank Ana M. Santander for expert assistance with protein purification, and Dr. Ramin Shiekhattar, Dr. Karim-Jean Armache, and members of their laboratories for helpful discussions.

We thank Dr. Haiyan Zheng and the Rutgers University Biological Mass Spectrometry Facility for mass spectrometry analysis, and Dr. Gary LeRoy at the Cancer Proteomics Shared Resource (CPSR; RRID: SCR_028491) of the Sylvester Comprehensive Cancer Center at the University of Miami for technical guidance on proteomics sample preparation and data interpretation. We also thank the Genome Technology Center at NYU Langone Health for high-throughput sequencing assistance. Computational analyses were supported by the High-Performance Computing (HPC) Facility of the Center for Health Informatics and Bioinformatics at NYU Langone Health and the Frost Institute for Data Science and Computing (IDSC) at the University of Miami.

Research reported in this publication was performed in part at the Onco-Genomics Shared Resource (OGSR) of the Sylvester Comprehensive Cancer Center at the University of Miami, RRID: SCR_022502, which is supported by the National Cancer Institute (NCI) of the National Institutes of Health (NIH) under award number P30CA240139.

This work was supported by grants from the National Institutes of Health (R01NS100897 and R01CA199652 to D.R.), the Howard Hughes Medical Institute (to D.R.), and the Sylvester Comprehensive Cancer Center at the University of Miami Miller School of Medicine (to D.R.).

## Author Contributions

**S.L.** and **D.R.** conceived the study. **S.L.** led the overall project, designed the experimental strategy, and performed the majority of experiments, including PCGF3 and FBRSL1 domain mapping, reporter assays, patient allele knock-in in hESCs, cortical neuronal differentiation, and ChIP-seq, ATAC-seq, and RNA-seq profiling. **S.L.** performed all bioinformatic and computational analyses, generated the figures, and wrote the manuscript with input from all authors. **H.-W.C.** performed FBRSL1 affinity purification and mass spectrometry, validated corepressor interactions and holocomplex assembly, conducted paralog displacement and reporter assays, and contributed to data interpretation. **D.R.** supervised the project, secured research funding, and edited the manuscript. All authors reviewed and approved the final manuscript.

---

**Figure S1.**
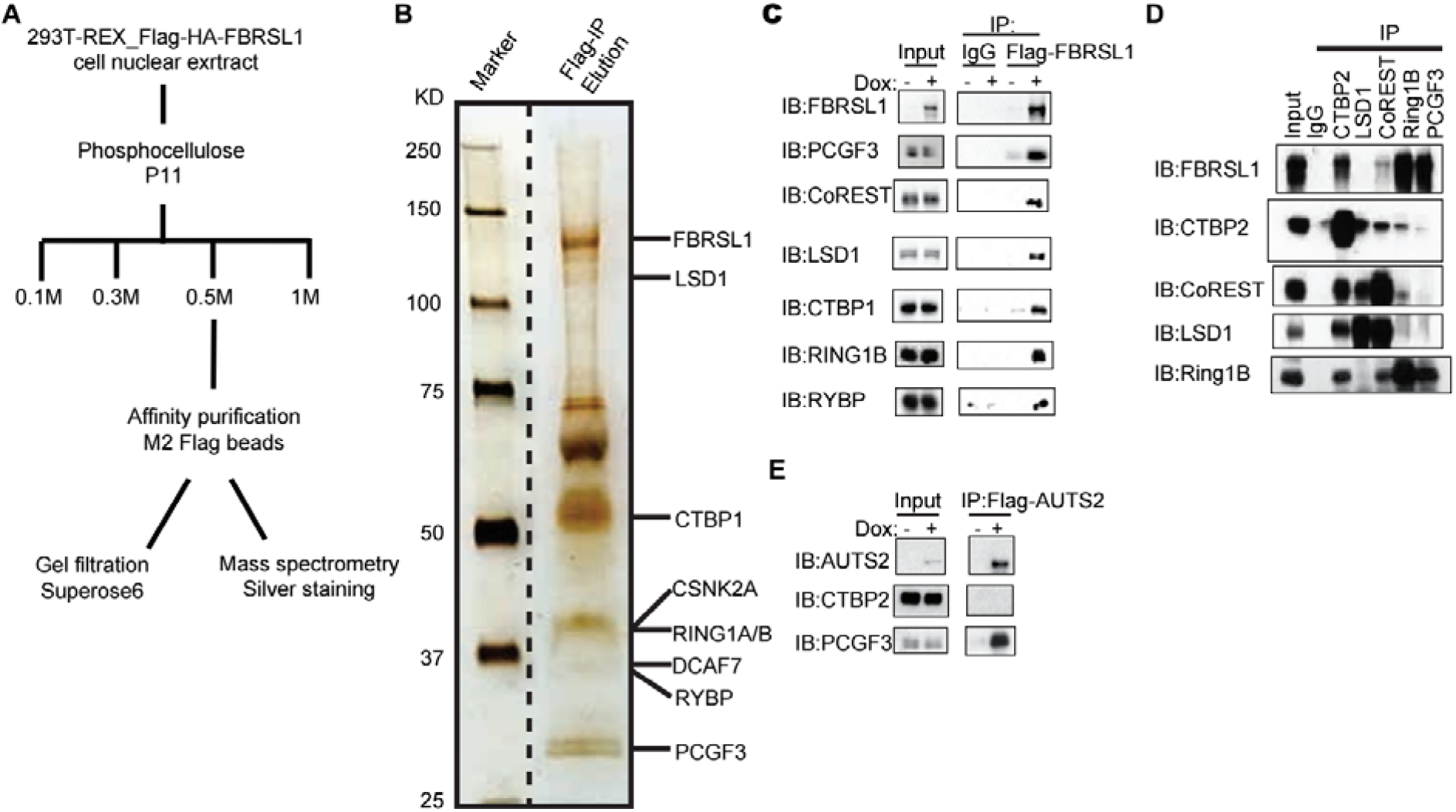
Biochemical validation of FBRSL1 association with ncPRC1.3 and CoREST/CTBP/LSD1 corepressors, related to Figures 1 and 2. **(A)** Scheme for purifying NFH-FBRSL1 from nuclear extract of HEK293T-REx cells with Dox-inducible Flag-HA-FBRSL1. Nuclear extract was fractionated on phosphocellulose P11, and the 0.5 M fraction was affinity-purified on anti-Flag M2 beads. The eluate was analyzed by silver staining and mass spectrometry (B; Figure 1D) or by Superose 6 gel filtration (Figure 2A). **(B)** Silver-stained SDS-PAGE of the Flag-peptide eluate from NFH-FBRSL1 purification. **(C)** Anti-Flag immunoprecipitation from NFH-FBRSL1 cells with or without Dox, with IgG as the control. Blots were probed for FBRSL1, the ncPRC1.3 subunits PCGF3, RING1B and RYBP, and the CoREST/CTBP/LSD1 complex subunits. **(D)** Reciprocal IP of endogenous CTBP2, LSD1, CoREST, RING1B and PCGF3, with IgG as the control, from HEK293T-REx nuclear extract. Blots were probed for FBRSL1, CTBP2, CoREST, LSD1 and RING1B. **(E)** Anti-Flag IP from NFH-AUTS2 cells with or without Dox. Blots were probed for AUTS2, CTBP2 and PCGF3.

**Figure S2.**
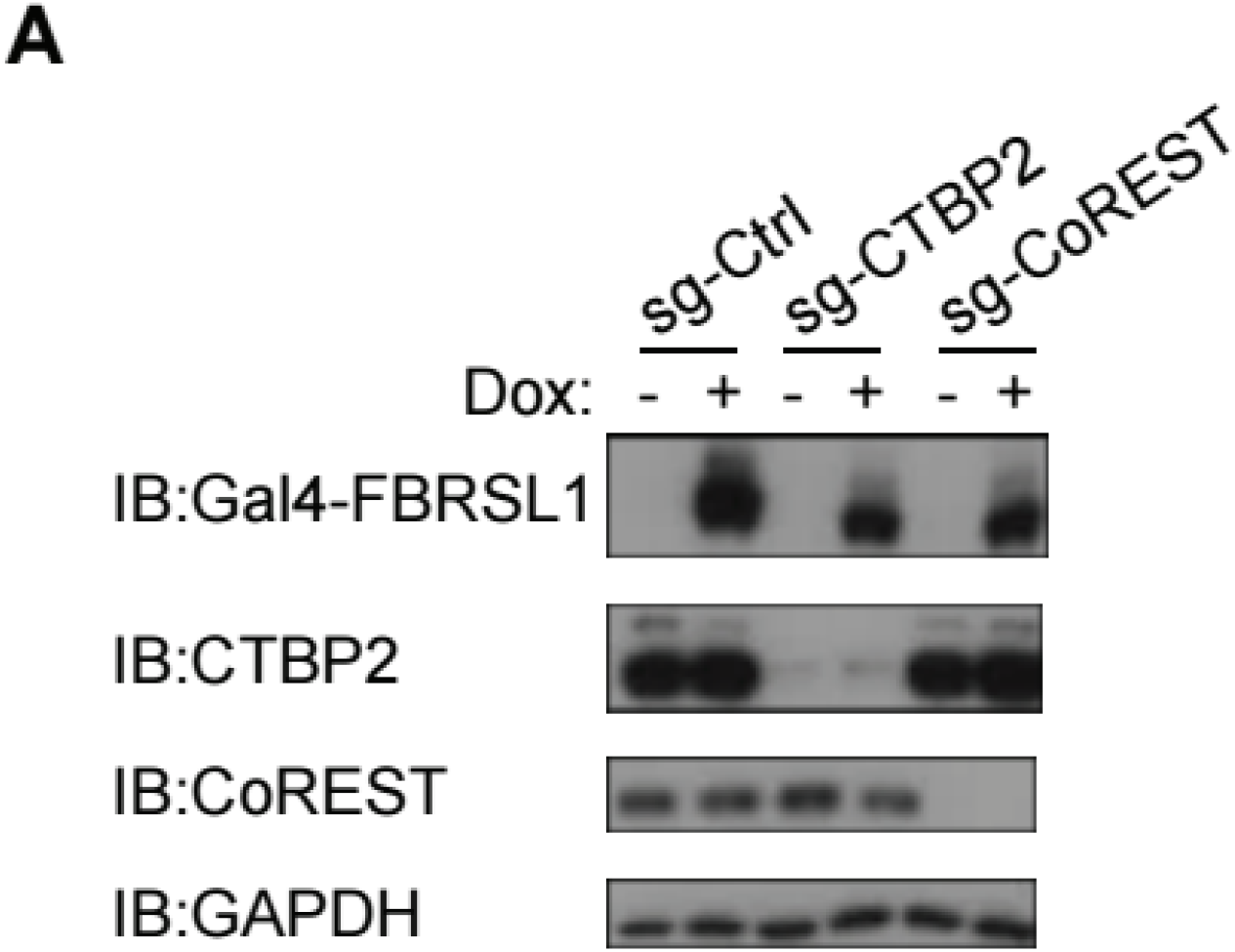
Validation of CoREST and CTBP2 KO and GAL4-FBRSL1 induction, related to Figure 2C. **(A)** Immunoblot of Gal4-FBRSL1 reporter cells expressing a control sgRNA (sg-Ctrl) or sgRNAs targeting CTBP2 or CoREST, with or without Dox. Blots were probed for Gal4-FBRSL1, CTBP2 and CoREST, with GAPDH as the loading control.

**Figure S3.**
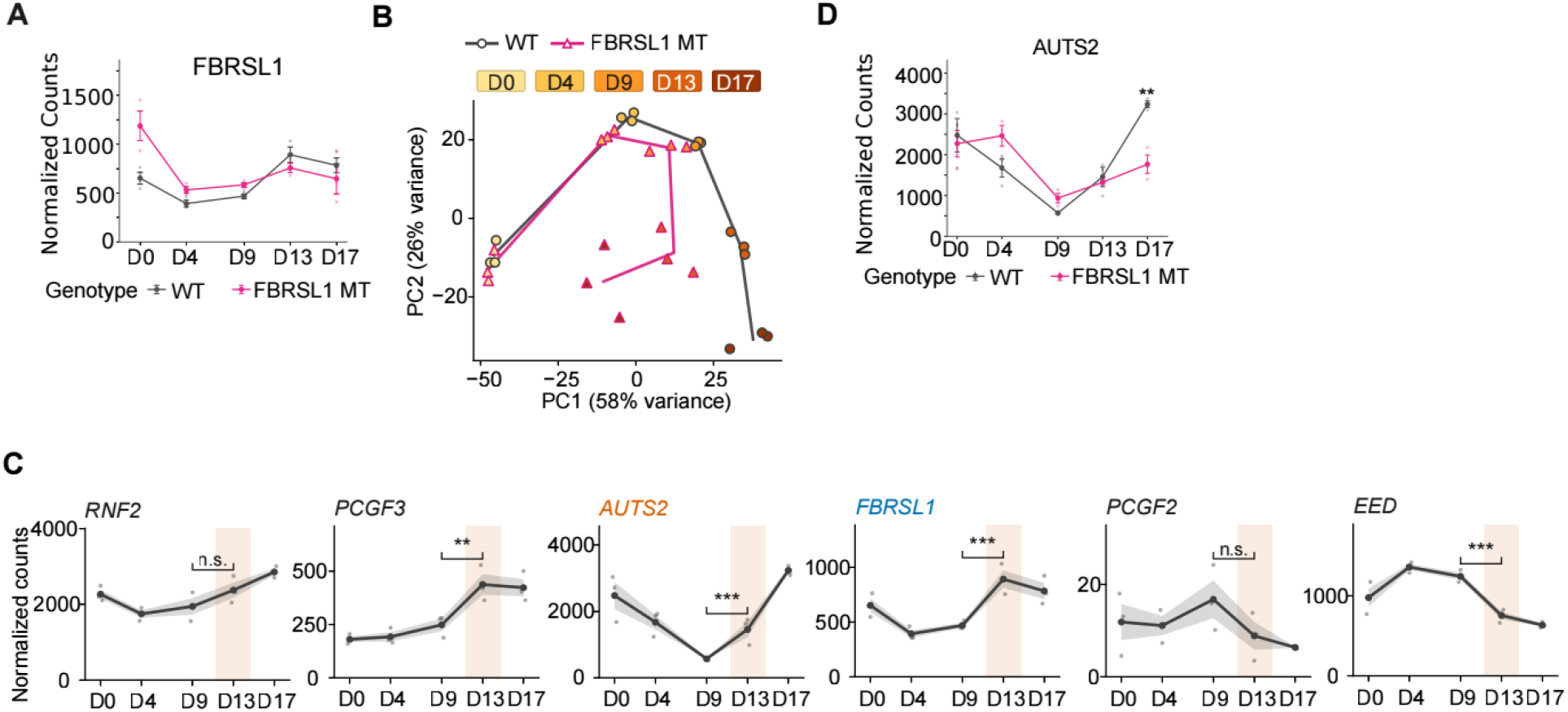
Expression trajectories of Polycomb subunits during cortical differentiation, related to Figures 5 and 6. **(A)** Expression trajectories of the *FBRSL1* in WT and FBRSL1-MT cells across differentiation. Values are DESeq2-normalized counts. Lines show the mean, and dots show individual replicates (n = 3). **(B)** Principal component analysis of all WT and homozygous FBRSL1 p.Gln163* (FBRSL1-MT) RNA-seq libraries using the 500 most variable genes after variance stabilization. Points represent individual libraries; color indicates day and shape indicates genotype. Lines connect day-specific genotype centroids. **(C)** Expression trajectories of *RNF2* (RING1B), *PCGF3*, *AUTS2*, *FBRSL1*, *PCGF2* and *EED* across WT differentiation. The shaded column marks D13. Brackets show the D13 versus D9 comparison (DESeq2 Wald test, Benjamini–Hochberg adjusted). **p < 0.01; ***p < 0.001; n.s., not significant. **(D)** Expression trajectories of the *AUTS2* in WT and FBRSL1-MT cells across differentiation. Values are DESeq2-normalized counts. Lines show the mean, and dots show individual replicates (n = 3).

**Figure S4.**
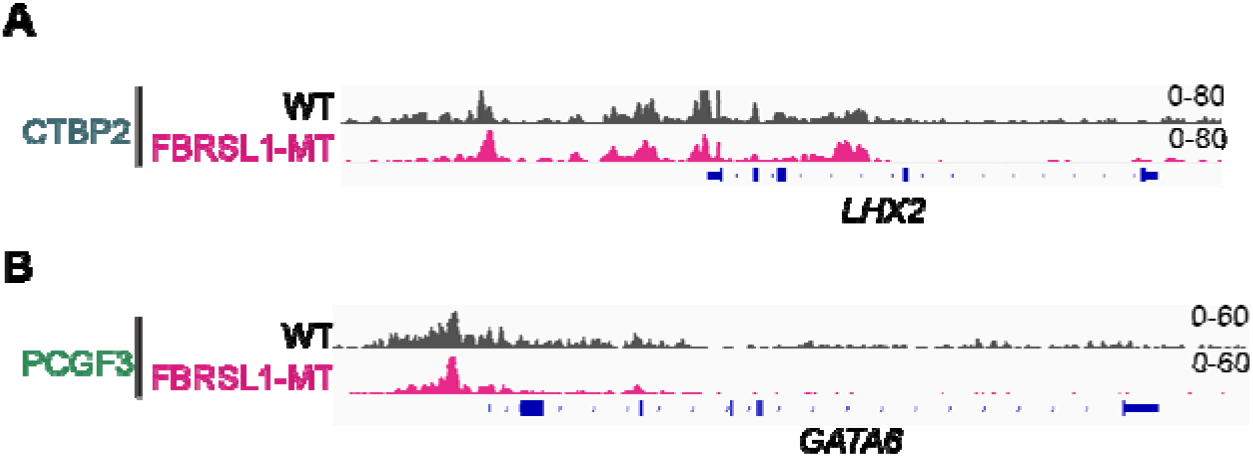
Retention of corepressor and Polycomb occupancy at control loci in *FBRSL1* mutant neural progenitors, related toy. **(A)** Genome browser tracks of CTBP2 ChIP-seq signal at the *LHX2* locus in wild-type (WT) and *FBRSL1* p.Gln163* neural progenitors at Day 13 (D13). **(B)** Genome browser tracks of PCGF3 ChIP-seq signal at the *GATA6* locus in WT and *FBRSL1* p.Gln163* neural progenitors at D13.

